# Response and oil degradation activities of a northeast Atlantic bacterial community to biogenic and synthetic surfactants

**DOI:** 10.1101/2020.12.18.423525

**Authors:** Christina Nikolova, Umer Zeeshan Ijaz, Clayton Magill, Sara Kleindienst, Samantha B. Joye, Tony Gutierrez

**Affiliations:** Institute of Mechanical, Process and Energy Engineering, School of Engineering and Physical Sciences, Heriot-Watt University, Edinburgh, EH14 4AS, United Kingdom; School of Engineering, University of Glasgow, Glasgow, G12 8LT, United Kingdom; Institute for GeoEnergy Engineering, School of Energy, Geoscience, Infrastructure and Society, The Lyell Centre, Edinburgh, EH14 4AS, United Kingdom; Center for Applied Geosciences, Eberhard Karls University of Tübingen, Tübingen, Germany; Department of Marine Sciences, The University of Georgia, Athens, GA, USA

**Author notes:** **Corresponding author:** Tony Gutierrez, Institute of Mechanical, Process and Energy Engineering, School of Engineering and Physical Sciences, Heriot-Watt University, Edinburgh EH14 4AS, UK.

**Keywords:** dispersant, biosurfactant, rhamnolipid, marine environment, crude oil, hydrocarbons, biodegradation, Faroe-Shetland Channel

## Abstract

**Background:** Although synthetic dispersants are effective in dispersing crude oil, they can alter the natural microbial response to oil and potentially hinder its biodegradation. Biosurfactants, however, are naturally derived products that play a similar role to synthetic dispersants in oil spill response but are easily biodegradable and less toxic. This study investigated the microbial community dynamics, ecological drivers, functional diversity, and oil biodegradation potential of a northeast Atlantic marine microbial community to crude oil when exposed to rhamnolipid or synthetic dispersant Finasol OSR52.

**Results:** We found the microbial community composition and diversity were markedly different in the rhamnolipid-amended treatment compared to that with Finasol, with key aromatic hydrocarbon-degrading bacteria like *Cycloclasticus* being suppressed in the Finasol treatment but not in oil-only and rhamnolipid-amended treatments. Psychrophilic *Colwellia* and *Oleispira* dominated the community in both the rhamnolipid and Finasol OSR52 treatments initially but later community structure across treatments diverged significantly: *Rhodobacteraceae* and *Vibrio* dominated the Finasol-amended treatment and *Colwellia, Oleispira*, and later *Cycloclasticus* and *Alcanivorax*, dominated the rhamnolipid-amended treatment. *Vibrio* abundance increased substantially in treatments receiving Finasol, suggesting a potentially important role for these organisms in degrading dispersant components. In fact, Finasol was linked with a negative impact on alpha diversity. Deterministic environmental filtering played a dominant role in regulating the community assembly in all treatments but was strongest in the dispersant-amended treatments. Rhamnolipid-amended and oil-only treatments had the highest functional diversity, however, the overall oil biodegradation was greater in the Finasol treatment, but aromatic biodegradation was highest in the rhamnolipid treatment.

**Conclusion:** Overall, the natural marine microbial community in the northeast Atlantic responded differently to crude oil dispersed with either synthetic or biogenic surfactants over time, but oil degradation was more enhanced by the synthetic dispersant. Collectively, our results advance the understanding of how rhamnolipid biosurfactants affect the natural marine microbial community, supporting their potential application in oil spills.

## Background

Extensive tracking of the microbial response to crude oil contamination in the ocean after the Deepwater Horizon oil spill in the Gulf of Mexico in 2010 provided an unprecedented view into feedbacks between environmental chemical signatures and microbial community evolution [1]. During this historic spill, approximately 700,000 tonnes (4.9 million barrels) of Louisiana light sweet crude oil was discharged into the Gulf from a blown-out wellhead at a depth of ∼1,500 m. Because of the scale and nature of the oil spill, synthetic dispersants were the primary response tool employed [2]. The decision to employ synthetic dispersants during a marine oil spill is driven largely by the desire to keep oil from reaching sensitive coastlines – often the primary goal of dispersant application. This unprecedented dispersant application involved approximately 7 million litres of the synthetic dispersants Corexit 9500 and 9527 to sea surface oil slicks and directly at the discharging wellhead along the seabed [3]. Prior to the Deepwater Horizon incident, limited knowledge of the effects of synthetic dispersants use on open ocean microbial communities was available. As a consequence, questions were raised about the response of autochthonous populations of hydrocarbon-degrading (hydrocarbonoclastic) bacteria – key players in oil biodegradation – to these dispersants and the need to identify the impact of dispersants on oil bioremediation was highlighted.

Following the Deepwater Horizon incident, a number of studies investigated the effects of Corexit on natural microbial communities; some studies also reported the response of oil biodegradation rates. Corexit appeared to inhibit natural microbial oil biodegradation in some cases, possibly due to the toxicity by one or more of the dispersant ingredients and/or because some microbes that responded to dispersants (e.g. *Colwellia* spp.) preferred to metabolize dispersant constituents more than oil [4, 5]. Some studies have reported that Corexit, and other synthetic dispersants, stimulated oil biodegradation by increasing its bioavailability to microorganisms [6–8]. Although the main components of synthetic dispersants are food-grade surfactants, including Tween 80 and Span 80, other components are hydrocarbon-based solvents that could confer toxicological impacts, whilst others are unknown as they are proprietary knowledge. Furthermore, a commonly used surfactant in dispersant formulations is dioctyl sodium sulfosuccinate (DOSS), a known toxin [9], that persists in the environment for months [10] to years in cold (deep sea) environments [11].

Hydrocarbonoclastic bacteria produce biosurfactants [12] that serve a similar purpose as synthetic dispersants, namely to reduce the surface and interfacial tension between oil droplets and seawater and increase the rate of oil biodegradation [12]. The most commonly studied biosurfactant producer is *Pseudomonas aeruginosa*, a ubiquitous bacterial species that grows on a wide range of hydrocarbon and non-hydrocarbon substrates and is known for its production of the glycolipid surfactant rhamnolipid with excellent surface-active properties (reduces the surface tension of water from 72 mN m^-1^ to less than 30 mN m^-1^ and facilitates formation of stable emulsions of petrol and diesel) [13]. Rhamnolipids have been shown to be effective in dispersing crude oil and enhancing its biodegradation by a pre-selected bacterial consortiums [14] some of which containing oil-degrading strains of *Ochrobactrum* sp. and *Brevibacillus* sp. [15]. However, studies comparing synthetic and bio-based surfactants effects on indigenous marine microbial communities are rare. We are aware of only one, albeit, recent study that compares the effects of a biosurfactant, in this case, a surfactin produced by *Bacillus* sp. strain H2O-1, to a synthetic dispersant, Ultrasperse II, with a natural marine microbial community, including its biodegradation of crude oil [16]. The surfactin enriched hydrocarbonoclastic bacteria more so than the synthetic dispersant, but no difference in oil biodegradation across treatments was observed.

We investigated the effect of rhamnolipid and the synthetic dispersant Finasol OSR52, which like Corexit, is stockpiled worldwide for use in oil spill response, on the composition of a natural marine microbial community and its ability to biodegrade crude oil. The study site was the Faroe-Shetland Channel (FSC), a subarctic region located on the UK Continental Shelf west of the Shetland Islands. The region is distinctive as it has a 20 year history of oil exploration and production, with some fields as deep as 1,500 m (e.g. Lagavulin) [17]. The FSC has complex and dynamic physical circulation characterized by mixing of distinct water masses [18] and the area is remote, cold, complex, and characterized by rough weather conditions for majority of the year, meaning that an oil spill response would be challenging. In addition, the FSC hosts important deep sea biological diversity, such as deep-sea sponges, cold-water coral communities, and a vibrant commercial fishing industry [18] which can be negatively impacted by major oil spills where synthetic surfactants were used.

Laboratory microcosm experiments in this study elucidated and compared the effects of the synthetic dispersant Finasol and the biosurfactant rhamnolipid on bacterial communities in the FSC and on their ability to biodegrade crude oil. A roller-table setup was employed to simulate sea surface conditions under *in situ* temperature, and crude oil was obtained from the Schiehallion oil field in the FSC (Fig. 1). Illumina MiSeq 16S rRNA sequencing was used to track microbial community abundance and dynamics, and the PICRUSt2 algorithm of QIIME2 was used to predict community functional diversity and abundance. Gas chromatography-Flame ionization detection coupled mass spectrometry (GC-FID/MS) was used to track the crude oil biodegradation in the microcosms.

**Figure 1.**
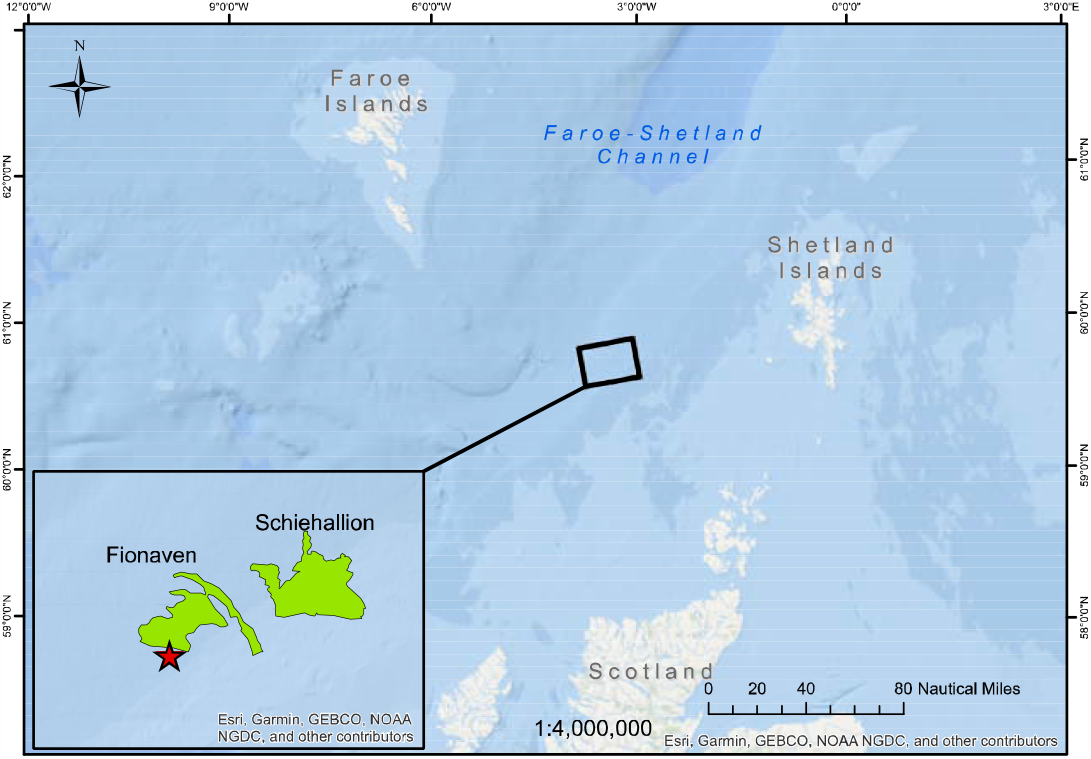
Map of the sampling site location (red star) and nearby oil-producing fields (green) in the Faroe-Shetland Channel. The map was created with ArcGIS Map software ver.10.6.1 (ESRI, USA) and freely available data from Oil & Gas UK.

## Materials and methods

### Field sampling and microcosm set-up

Surface seawater was collected on May 2018 from 3 m depth in the Faroe-Shetland Channel (FSC) (60°16.36’ N, 04°20.60’ W; Figure 1), which is a subarctic, deep-water region of the northeast Atlantic characterised by an active oil and gas industry (Supplementary Methods). Immediately after sampling, the seawater was transferred onboard to 10 L carboys and stored at 4°C and used within 2 days in the set-up of the various experiments after returning to the laboratory at Heriot-Watt University.

To assess the changes in the microbial community structure and dynamics during enrichment with crude oil and in the presence of either the synthetic dispersant Finasol, or the biosurfactant rhamnolipid, seven microcosm treatments were set up with aliquots from the WAF mixtures as described in detail in Supplementary Information. Briefly, five microcosm treatments (WAF, CEWAF, BEWAF, SWD, and SWBS) and untreated controls were prepared using 0.5-L glass bottles (each containing 300 ml of sample volume) and incubated on a roller table for 4 weeks. Sampling was performed at the beginning of these incubations (day 0) and then subsequently thereafter at days 3, 7, 14 and 28. The WAF, CEWAF, BEWAF, SWD, and SWBS were prepared by mixing filter-sterilized seawater with oil and/or the synthetic dispersant (Finasol) or the biosurfactant (rhamnolipid) for 48h at room temperature and subsequently the dissolved fraction, excluding contamination by oil or dispersant/biosurfactant. To analyse for changes in the hydrocarbon composition of the oil due to biodegradation, replicates of the treatments that contained the oil (i.e. the WAF, BEWAF and CEWAF treatments) were prepared (in triplicate) in identically the same way as described above (Supplementary Methods).

### DNA extraction and barcoded amplicon sequencing

DNA was extracted according to the method of a previous study [19] which utilizes chemical cell lysis with potassium xanthogenate buffer. DNA extracts were resuspended in 20 μl of 1 mM TE buffer and stored at -20°C for Illumina barcoded-amplicon sequencing. A two-step amplification procedure (Supplementary Methods) was used to amplify the 16S rRNA gene in order to minimize heteroduplex formation in mixed-template reactions [20]. The purified PCR reactions were pooled together and then sent for paired-end Illumina MiSeq sequencing (Illumina 2 × 250 v2 kit) at the Edinburgh Genomics Facility (University of Edinburgh, UK).

### Bioinformatic and statistical analysis

The resulting 16S rRNA gene sequences were processed with the open-source bioinformatics pipeline QIIME2 [21]. Initially, sequences were demultiplexed and quality-filtered using the DADA2 algorithm as a QIIME plugin [22]. DADA2 implements a quality-aware correcting model on amplicon data that denoises, removes chimeras and residual PhiX reads, dereplicate DNA reads, and calls amplicon sequence variants (ASVs) [23]. The quality-filtered sequences were then aligned to the reference alignment database SILVA SSU Ref NR release v132 [24]. 16S rRNA gene sequencing resulted in 7 703 409 pair-end sequence reads across 90 samples.

PICRUSt2 algorithm as a QIIME plugin [25] was used on the 16S rRNA gene sequences to predict the functional abundance and diversity of the microbial community (Supplementary Information). Whilst metabolic potential from 16S rRNA studies are often discounted as mere predictions, with the newer version of PICRUSt2 with a comprehensive reference database (>20,000 genomes covered as opposed to its predecessor which had ∼2000 only), and with a very high correlation with matched metagenomics datasets (∼0.9), and the fact that majority of the ASVs in our datasets were represented in the reference database, this analysis has a very strong utility to give mechanistic understanding.

Statistical analyses of the ASVs (alpha and beta diversity, PERMANOVA, PCoA, subset and regression analyses, functional abundance and diversity) were performed using the statistical software programme R-Studio v3.6.3 [26] and further described in detail in Supplementary Methods. The R scripts used to generate the analyses are available at http://userweb.eng.gla.ac.uk/umer.ijaz/bioinformatics/ecological.html and as part of R’s microbiomeSeq package http://www.github.com/umerijaz/microbiomeSeq.

## Results

### Bacterial community composition and diversity

Members of the Proteobacteria dominated the taxonomic profiles over the 28-day microcosm incubations, ranging between 60-98%. In contrast, the Bacteroidetes decreased from ∼40% (initially) to almost undetectable by the end of the experiment. Other phyla were present at <5% relative abundance. At day 0, all treatments showed similar community composition, comparable to the *in-situ* community. Community profiles were dominated at the family level by *Colwelliaceae, Saccharospirillaceae, Rhodobacteracea* and *Micavibrionaceae* (>40%; Fig. 2, Supplementary File 2).

**Figure 2.**
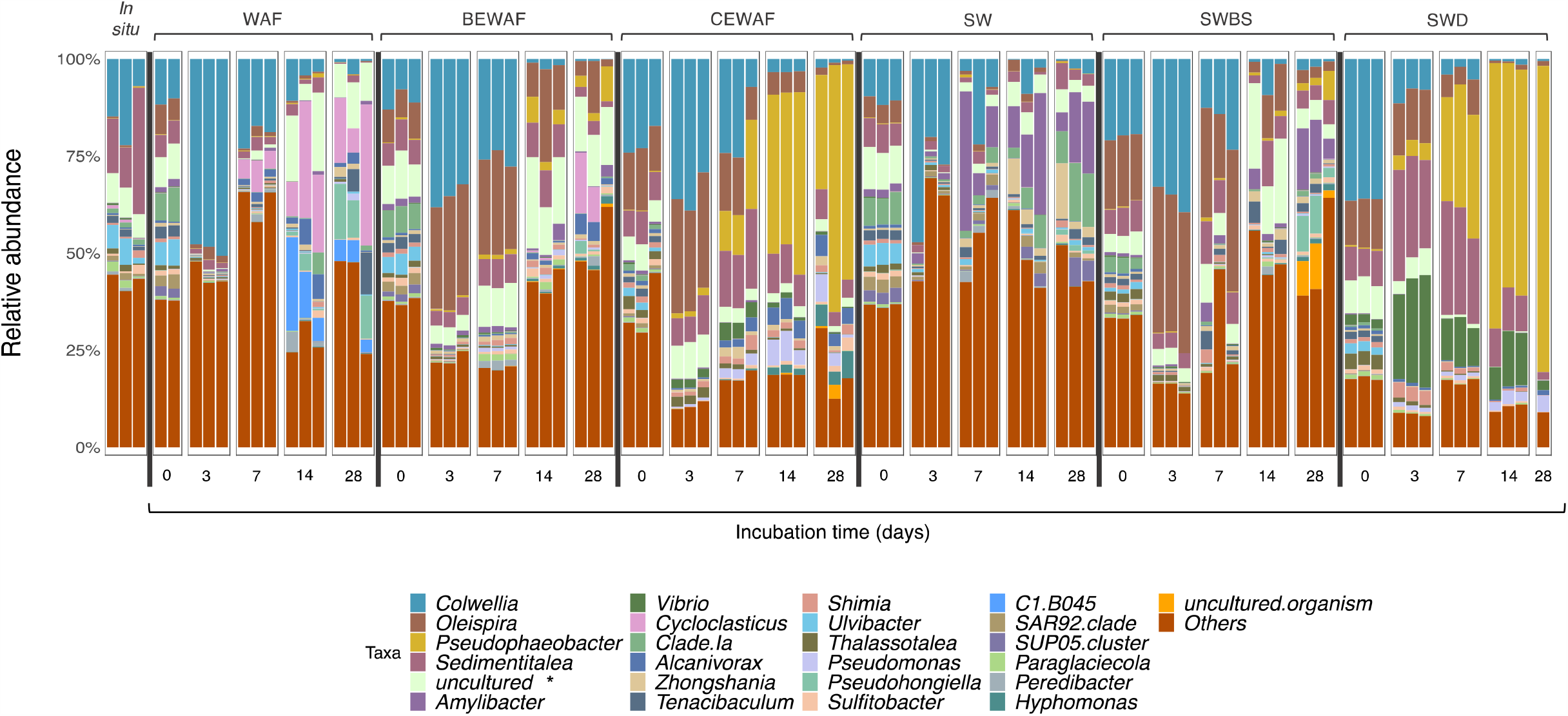
Relative abundance of top 25 most abundant taxa shown to genus level. Treatments at different incubation times are shown as independent triplicates where In situ is baseline microbial community at time of seawater sampling (FSC), WAF - seawater and oil only, BEWAF – crude oil and biosurfactant, CEWAF – crude oil and synthetic dispersant, SW - seawater only, SWBS - seawater and biosurfactant, and SWD – seawater and synthetic dispersant. *Represents uncultured bacteria from the *Micavibrionaceae* family. SWD had one replicate on day 28 and WAF had two replicates on day 0.

The initial abundance of *Colwellia* varied significantly across treatments. At day 0, *Colwellia* abundance ranged from 15% in the seawater control (SW) to 11% (WAF, BEWAF) to >35% in the dispersant only control (SWD) and CEWAF treatments. In the dispersant only control treatment (SWD), the abundance of *Colwellia* rapidly decreased to 9%, then to 5%, and then 1% on days 3, 7, and 28 respectively. The abundance of *Oleispira* increased to 28% in BEWAF and CEWAF treatments but was negligibly abundant in the oil-only treatment (Fig. 2, Supplementary File 2). By day 7, members of uncultured *Micavibrionaceae* increased in both the BEWAF and WAF treatments, peaking on day 14 (18% and 14%, respectively). On day 14 the community profiles of the WAF and BEWAF treatments were quite distinct, with *Colwellia* having markedly decreased in abundance in both treatments to 6% and 2%, respectively. Across all treatments *Cycloclasticus* and *Alcanivorax* were rare initially (<1%) but had increased by up to 20% by day 14 in the WAF treatment and these abundances were maintained on day 28. In the BEWAF, *Cycloclasticus* and *Alcanivorax* abundances remained below 1% until day 14 but increased to 8% and 4% by day 28, respectively. They were not detected in the seawater only control (SW). Oil degraders, except *Colwellia*, were not enriched in the SW treatment. Similar to the BEWAF and WAF treatments, on day 3 the CEWAF treatment was dominated by *Colwellia* (35%), *Oleispira* (28%), *Sedimentitalea* (8%) and uncultured members of the *Micavibrionaceae* (9%). *Pseudophaeobacter* and *Sedimentitalea* (family *Rhodobacteracea*) were exclusively enriched in the Finasol-ammended treatements (CEWAF and SWD) by the end of incubation. *Alcanivorax* increased in the CEWAF treatment from <1% in the early stages of incubation to 5% by the end.

A strong enrichment in *Vibrio* was observed only in the SWD treatment, with abundance increasing from 2% at day 0 to 26% by day 3, followed by a gradual decrease to 12% (day 14), and then to 2.4% (day 28). In the CEWAF treatment, for comparison, *Vibrio* became only slightly enriched (1 - 4%) throughout the incubation period. *Pseudomonas* was observed mainly in the CEWAF and SWD treatments where its abundance increased from <1% on day 3, to 6% on day 14 in the CEWAF treatment, and 3% in SWD. In the CEWAF treatment, *Cycloclasticus* was absent.

A phylogenetic tree of top 50 ASVs was constructed with their abundance changes shown for each treatment (Fig 3). In the top 50 ASVs, six ASVs were assigned to *Colwellia*, four to *Oleispira*, three to *Cycloclasticus*, one each to *Alcanivorax, Pseudomonas, Zhongshania, Thalassotalea, Shimia, Sulfitobacter*, and *Amylibacter*. Three of the *Colwellia* ASVs displayed higher abundances in the CEWAF treatment, while the other three distinct ASVs were enriched in the WAF. *Cyclocalsticus* ASVs were enriched exclusively in the WAF treatment, and *Alcanivorax* dominated in all the oil-amended treatments (WAF, BEWAF, CEWAF). Although *Pseudomonas* had overall low abundance, it was enriched in the CEWAF and SWD treatments.

**Figure 3.**
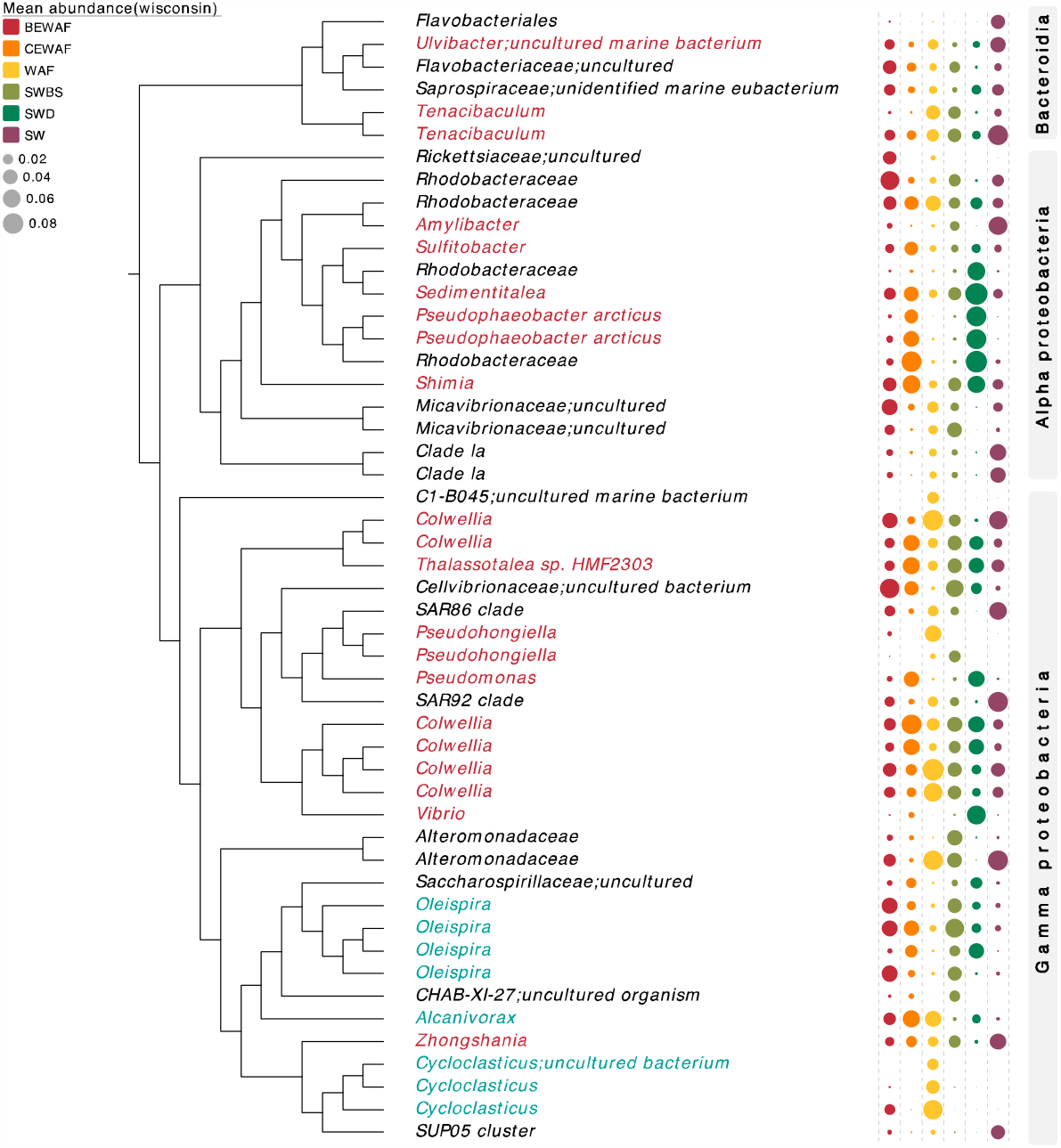
Phylogenetic tree showing the top 50 most abundant ASVs which have been taxonomically assigned with SILVA SSU v132 database. ASVs in red font represent known generalist hydrocarbonoclastic bacteria, and ASVs in blue font represent obligate hydrocarbonoclastic bacteria. Dot plots on the right side of the tree show the mean abundance of each ASV coloured by treatment after performing proportional standardization using the Wisconsin function.

### Bacterial diversity

Alpha diversity indices, species richness and Shannon index, were calculated across treatments and time. Species richness was highest in the BEWAF treatments and significantly different from Finasol-amended treatments which had the lowest diversity (Fig. 4A). Temporal changes across treatments revealed that the richness was the lowest on day 3 and after that gradually increased for all treatments (Supplementary Fig. S1), albeit with some small differences. Principle coordinates analysis (PCoA) plot revealed distinct clustering of treatments based on Bray-Curtis dissimilarities (Fig. 4C). Taking into account weighted UniFrac distance measure, all treatments clustered close to each other and even overlapped, indicating diversity similarity between the treatments. Treatment and incubation time were significant factors affecting the beta diversity (PERMANOVA, p=0.001) with treatment explaining up to 45% of the variability and time up to 26% (Figure 4C). The diversity variation patterns across treatments on a temporal scale are further discussed in Supplementary Results (Supplementary File 4). Overall, PERMANOVA showed that treatment and incubation time significantly influenced the variability in microbiome structure. Next, we performed subset regression analysis on one-dimensional realisation of the microbiome (alpha and beta diversity) to further understand which of treatments and/or incubation time points specifically caused an increase or decrease in the microbiome properties. The subset regressions confirmed that the presence of Finasol in the treatments decreased richness and Shannon entropy but increased the beta diversity (i.e. caused distinct clustering of samples). In contrast, the inclusion of rhamnolipid led to increase only in Shannon entropy and NRI and had an insignificant influence on the rest of the diversity measures (Supplementary Fig. S3). Further interpretation of the regression analysis is provided in Supplementary Results.

**Figure 4.**
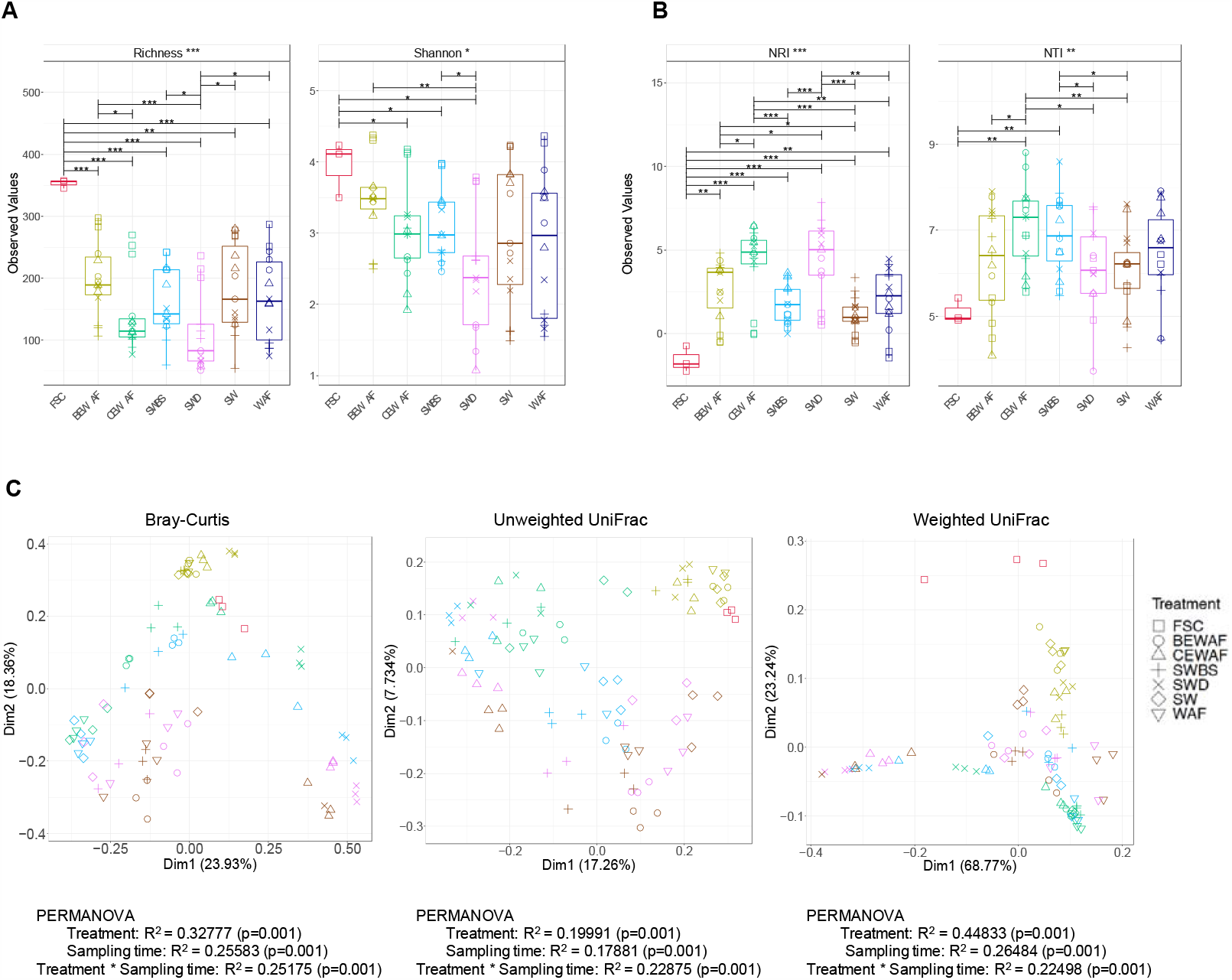
**(A)** Overall alpha diversity indices of ASVs and **(B)** net relatedness index (NRI) and nearest-taxon index (NTI). Statistically different treatments (pair-wise ANOVA) are connected by bracket and the level of significance is shown with: * (p<0.05), ** (p<0.01), or *** (p<0.001). Colours represent treatments and shapes the incubation time (square – day 0, plus – day 3, cross – day 7, circle – day 14, and triangle – day 28). **(C)** Principal Coordinate Analysis (PCoA) using Bray-Curtis, Unweighted Unifrac and Weighted Unifrac distance matrices. Ellipses represent 95% confidence interval of the standard error of the ordination points of a given grouping. Results from PERMANOVA test for each distance matrix are shown underneath each plot. Colours in **(B)** represent sampling time (red – in-situ seawater at time of collection, olive green – day 0, green – day 3, blue – day 7, pink – day 14, and brown – day 28).

In order to identify key taxa representing major shifts in the communities across the different treatments, we performed differential abundance analysis with DESeq2 with adjusted p-value significance cut-off of 0.05 and log2 fold chance, and the results described and presented in detail in the Supplementary Results and Supplementary File 5. Next, we considered subset analysis where we determined the minimum set of significant ASVs directly responsible for driving the community dynamics for each treatment over time. The subset analysis procedure calculates pair-wise Bray-Curtis distances between samples using all the ASVs in the abundance table, then permutes through the combination of ASVs until a minimal subset of ASVs, which beta diversity is conserved against the full ASV table by maximising the correlation, is found. We used differential heat trees to showcase how members of the highest correlated subsets for each treatment changed their abundances over time. The resulting reduced-order subsets correlated highly with the full table by preserving the beta diversity between samples (Fig. 5). In the BEWAF treatment, *Alteromonadaceae, Pseudophaeobacter*, members of the *Rhodobacteraceae* family (*Amylibacter* and *Sedimentitalea*), *Colwellia, Oleispira* and *Micavibrionaceae* significantly drove the shifts in community dynamics over time (PERMANOVA R^2^ = 0.395, p = 0.001), similarly to the seawater control (SW) (Supplementary Fig. S4). In the WAF treatment, the taxa driving the observed shifts in community diversity over time (PERMANOVA, R^2^ = 0.262, p = 0.007) are *Oleispira* and *Alcanivorax*, and putative oil-degraders *Colwellia* and *Pseudophaeobacter*. In the CEWAF treatment, only two taxa, unclassified members of the *Alteromonadaceae* family and *Amylibacter*, drove the diversity shift with 0.96 correlation with the full ASV abundance data (PERMANOVA, R^2^ = 0.539, p = 0.001).

**Figure 5.**
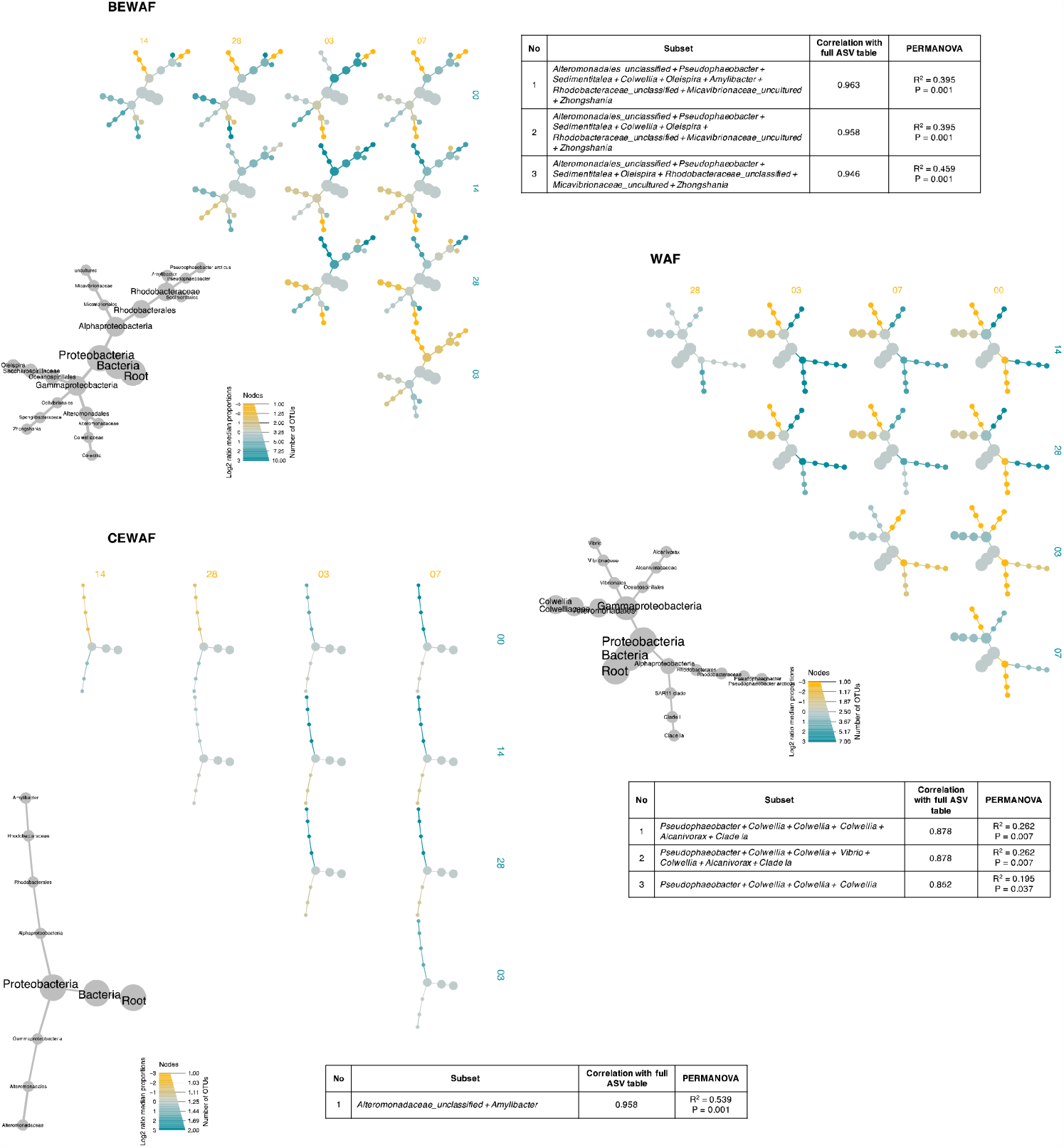
Differential heat trees showing the key differential taxa (DESeq2; using Wilcoxon p-value test adjusted with multiple comparison) in treatments BEWAF, WAF, and CEWAF. Subset analysis was performed to identify the subset of significant ASVs causing the major change in beta diversity in these treatments. The top 3 (where possible) subsets with the highest correlation with the full ASV table considering Bray-Curtis distance (PERMANOVA) are listed for each treatment. The grey trees are taxonomic key for the smaller unlabelled coloured trees. The colour of each taxon represents the log-10 ratio of median proportions of reads observed in each treatment. The size of tree nodes shows the number of ASVs (note: labelled as OTUs) present in each treatment.

### Ecological drivers of microbial assembly

Ecological processes responsible for changes in microbial communities in each treatment over time were determined by NTI and NRI which reveal whether stochastic (predation, competition, unpredicted disturbance etc.) or deterministic (selection imposed by an external environmental factor, i.e. environmental filtering). All treatments had NTI values were significantly greater than +2 (ANOVA, p<0.05), indicating strong clustering driven by deterministic environmental filtering (Fig. 4B). the addition of crude oil, either by itself or in combination with Finasol or rhamnolipid, determined the microbial community structure promoting co-existence of closely related and ecologically similar taxa. The NRI reflects the phylogenetic clustering in a broad sense (entire phylogenetic tree) with the negative values representing evenly spread communities, whereas values above 0 represent clustered communities. As expected, the environmental setting had no effect on the community structure in the *in situ* FSC community as indicated by negative NRI value (Figure 4B; Supplementary Fig. S5). The relative influence of environmental filtering on the microbial community composition varied over time but was strongest in the Finasol-amended treatments (CEWAF and SWD). At the beginning of the incubation all treatments had NRI values ∼0 suggesting that phylogenetic overdispersion (i.e. stochastic process) dominated the assembly processes in those communities. Over time, as NRI values increased, environmental pressure (*i*.*e*., the imposed treatment regime) determined the response of the bacterial community rather than stochastic processes.

### Functional diversity and abundance

The ASV sequences were further used to characterise the functional diversity of KEGG orthologs (PICRUSt2) as alpha and beta diversity. The predicted functional alpha diversity was calculated using two standard ecological diversity measures – the rarefied genetic functional richness and the Shannon entropy. Pairwise ANOVA on richness and Shannon indicated significant differences in the different treatments (Supplementary Fig. S6). Treatments at each incubation time points were also compared to identify significant changes over time (Supplementary Fig. S7A). The predicted richness of KO was the highest in the BEWAF treatment, and statistically different from the CEWAF and SWD treatments on days 3 and 7 compared to the rest of the treatments during the same period (Supplementary Fig. S7A). By days 14 and 28, the richness in the BEWAF and WAF treatments was very similar and close to the maximum number of predicted KO. The SWD treatment displayed the lowest diversity of KO (Supplementary Fig. S6), especially on day 7 and thereafter (Supplementary Fig. S7A). The beta diversity was calculated by Bray-Curtis dissimilarity distance which demonstrated a distinct clustering of all treatments on day 0, and for SWD on days 3, 7, and 14, while the rest of the treatments overlapped (Supplementary Fig. S7B). PERMANOVA analysis revealed that the treatment and incubation time were significant factors (PERMANOVA, p=0.001) and which respectively explained 18% (R^2^ = 0.1814) and 21% (R^2^ = 0.2064) of the variability in beta diversity of KEGG orthologs. A list of all significantly different pathways between all of the treatments is presented in Table S12, where KO hits for each pathway were presented.

To present the predicted functional abundance, we selected specific KO involved in aliphatic and aromatic hydrocarbon degradation pathways, as well as biosurfactant synthesis. For simplicity, we examined only the three treatments containing crude oil – BEWAF, CEWAF and WAF. Samples from CEWAF treatment showed potential enrichment of genes involved in the degradation of medium-chain length alkanes, styrene, fluorobenzoate, PAHs (naphthalene, phenanthrene etc.), chlorocyclohexane, chlorobenzene, and xylene (Supplementary Fig. S8). The relative abundance of genes which encode for the degradation of short-chain length (methane monooxygenase) and medium-chain length (alkane 1-monooxygenase and rubredoxin-NAD(+) reductase) alkanes were significantly increased in all three treatments on day 3, and in the CEWAF on days 3 and 7. The BEWAF treatment was enriched with genes involved in the degradation of chloroalkanes, benzoate, bisphenol, furfural, and flurobenzoate, whilst genes involved in the degradation of BTEX (benzene, toluene, ethylbenzene, xylene), dioxin and nitrotoluene were predicted to be more abundant in the WAF treatment (Supplementary Fig. S8).

Genes for different biosurfactants were also predicted. Genes involved in rhamnolipid synthesis, namely *rhl*A and *rhl*B (rhamnosyltransferases) were predicted at days 0 and 3 in the BEWAF, CEWAF and WAF treatments, although their abundance was not high (Supplementary Fig. S8). Surfactin synthesis genes were also predicted in these same treatments and time period, but with relatively higher abundance. Exopolysaccharide production protein ExoY was detected in all the treatments, but at lower abundance than for the rhamnolipid and surfactin genes on days 0 and 3, but it was comparatively higher on day 14 in the BEWAF treatment.

### Hydrocarbon biodegradation

The GC-FID chromatograms for each oil-amended treatment were compared in order to assess the extent of degradation over the course of the incubation (Fig. 6). The peak areas for C_12_ to C_30_ *n*-alkanes, and two PAHs, phenanthrene and methylphenanthrene were used to calculate ratios of specific hydrocarbons indicative of biodegradation (Supplementary Fig. S9). The oil biodegradation in the rhamnolipid-amended treatments was relatively slow but insignificant (ANOVA, p>0.05) in the first week of incubation but it was significantly faster (ANOVA, p<0.05) between day 14 and day 28 as indicated by pristane/*n*C_17_ ratio (Fig. 6). In contrast, oil biodegradation in the Finasol-amended treatment was initially rapid but slowed down over time, whereas biodegradation of alkanes in the WAF treatment was insignificant over time. PAH degradation was not significantly different across treatments or sampling times, though concentrations generally decreased over time in all treatments. The phenanthrene/9-methylphenanthrene ratio in the BEWAF treatment decreased the most by the end of the incubation compared to that in the CEWAF and WAF treatments (Fig. 6; Supplementary Fig. S9), suggesting the highest rates of PAH degradation in the BEWAF treatment.

**Figure 6.**
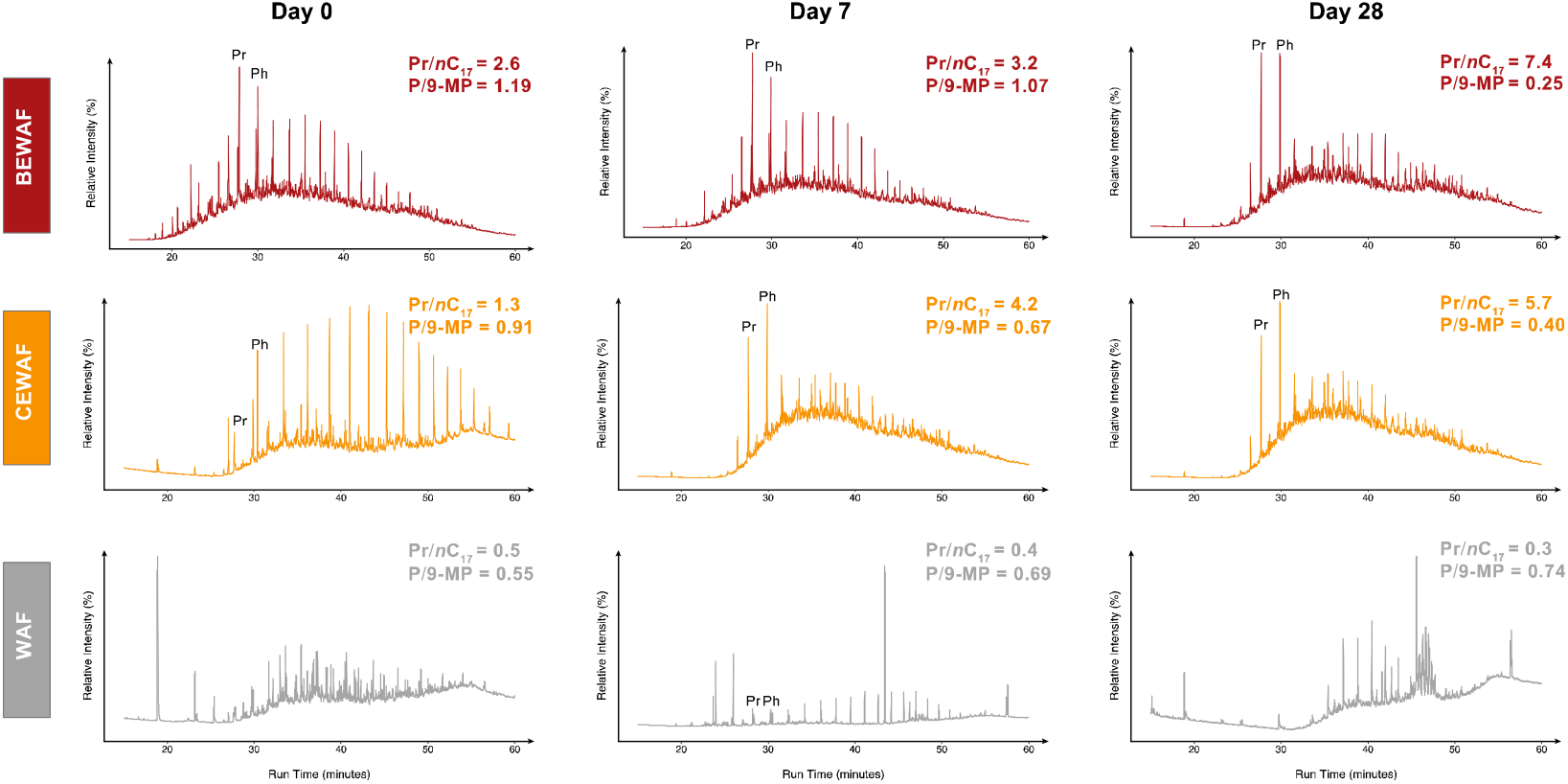
Representative flame-ionization chromatograms of the aliphatic hydrocarbon fraction of BEWAF (red), CEWAF (orange), and WAF (grey) through time of incubation (days 0, 7 and 28). Also shown are ratios of pristane versus heptadecane (Pr/*n*C17), which increases with increased biodegradation, and phenanthrene versus 9-methylphenanthrene (P/9-MP), which has an inverse relationship with biodegradation extent. Pristane (Pr) and phytane (Ph) are annotated for reference. Note ordinate axis is displayed in relative abundance.

## Discussion

This study probed the response of bacterial communities in surface waters of the FSC to crude oil alone or in concert with a biosurfactant (rhamnolipid) or a government-approved synthetic dispersant, Finasol OSR52. Most studies describing the microbial response to dispersed crude oil have focused on the effect of synthetic dispersants. The effectiveness of biosurfactants for dispersing oil and enhancing its biodegradation is well known [27, 28], but they have not been compared with synthetic dispersants directly to date. No published studies have investigated the effect of rhamnolipids on the microbial community response to crude oil in cold marine environments under simulated *in situ* conditions. Since a number of studies after the Deepwater Horizon spill documented negative impacts of synthetic dispersants [3–5] the need to explore alternative products, such as biosurfactants, and to assess microbial responses under environmentally-relevant conditions is critical [2] The potential for developing microbial biosurfactants for combatting oil spills is fertile territory that has been scarcely explored.

### Bacterial community dynamics

The FSC is a cold subarctic environment (avg. T=9.7°C), so *in situ* bacterial communities were dominated by psychrophilic taxa, including known oil-degraders, *e*.*g*., *Oleispira, Colwellia* and *Cycloclasticus*, that reside in cold sea surface [8, 29–31] and subsurface [32] waters, including the FSC [33, 34]. The high abundance of oil-degrading bacteria and their rapid (within 3 days) response to the crude oil, suggests they are primed from background exposure to hydrocarbons, possibly through permitted releases of produced water or from adjacent North Sea waterways Frequent shipping and oil transportation activities in and around the FSC, including from the Sullom Voe, a major terminal on the west coast of Shetland, may also generate hydrocarbon exposure. Though no confirmed natural oil seeps are known in the FSC or nearby, evidence from satellite surveys showing oil slicks suggest subsurface oil seeps on the east and west of Scotland and offshore in the North Sea (Peter Browning-Stamp, pers. comm.). Natural seepage is known to prime a rich community of oil-degrading bacteria in the Gulf of Mexico [1] and the FSC appears to behave similarly.

*Colwellia* are commonly observed in cold surface and deep sea environments [31, 33, 35], and some members of the genus utilise a broad range of hydrocarbons, including short-chain alkanes [36], as well as PAHs, *e*.*g*., phenanthrene [37]. The metabolic versatility of *Colwellia* likely explained its early bloom in the WAF, BEWAF and CEWAF treatments but this may also result from their high *in situ* abundance in the FSC. In a recent study, *Colwellia* were implicated in dispersant-component degradation in treatments of deep-sea water from the Gulf of Mexico amended with the dispersant Corexit; their abundance increased from 1% to 43% after only one week [4]. Interestingly, in this study, *Colwellia* was the dominant organisms in the CEWAF treatment (by day 3), but in the dispersant-only treatment (SWD) the abundance of these organisms decreased markedly by day 3, and continued to decline thereafter subsequently becoming overprinted by members of the *Rhodobacteracaea* and *Vibrionaceae*; this pattern was similarly reported in another study using the synthetic dispersant Superdispersant-25 [33].

Members of the *Rhodobacteracaea* and *Vibrionaceae* may utilize oil-derived organic intermediates produced by hydrocarbon degraders [38, 39] or they may consume components of the dispersant itself, as previously shown in other studies using Corexit [4]. *Vibrio* became markedly enriched in only the SWD treatment, with highest levels reached by day 3. In contrast, *Vibrio* was less abundant in the CEWAF treatments, suggesting that these organisms might have a preference for the Finasol over the oil as a carbon and energy source. *Vibrio* are known for their quorum sensing ability and it is possible that their metabolic agility [40] allowed them to outcompete *Colwellia* in the SWD treatment. The observed bloom of the *Rhodobacteracaea*, mainly *Sedimentitalea, Pseudophaeobacter* and to a lesser extent *Sulfitobacter*, may have contributed to oil degradation as members of these genera have hydrocarbon degrading capabilities [34, 39].

A previous study [4] employed the dispersant Corexit 9500, whose composition (18% DOSS, 4.4% Span 80, 18% Tween 80, and 4.6% Tween 85) is distinct from Finasol (15-25% DOSS, 15-23% non-ionic carboxylic acids and alcohols). The selection of *Colwellia* could be driven by DOSS, the common agent in both dispersants. However, the presence of other surfactants in Finasol may have promoted a sustained response by *Vibrio*, especially in the dispersant only (SWD) treatment. A previous study (38) found that *Vibrio* responded strongly to Corexit amendment, mirroring observations of increased *Vibrio* abundance in impacted Gulf of Mexico surface water samples collected during the active discharge phase of the incident [41]. Techtmann et al. [42] suggested that *Vibrio* metabolized metabolic by-products of oil degradation by other microorganisms. This could not be the mechanism in this case, however, since there is no oil in the SWD treatment, but it is possible that the *Vibrio* were metabolizing the carboxylic acids and alcohols in the dispersants. Similar compounds are known intermediates of oil biodegradation [43].

*Alcanivorax* was only observed in oil-amended treatments; it was undetectable in the controls (SW, SWD and SWBS). *Alcanivorax* is recognized for its almost exclusive preference for aliphatic hydrocarbons and for commonly blooming soon after oil is introduced to an environment [12, 30, 33]. However, *Alcanivorax* increased later (days 14 and 28) in the WAF, BEWAF, and CEWAF treatments. This was unexpected and may be because they were outcompeted by more resilient earlier bloomers (esp. *Colwellia, Oleispira, Sedimentitalea*, and uncultured *Micavibrionaceae*), which is reminiscent to the non-enrichment of *Alcanivorax* during the Deepwater Horizon oil spill. 16S rRNA gene sequences for this genus were undetected in water column metagenomic libraries from the Gulf of Mexico during the spill’s most active phase [35, 37]. *Alcanivorax* in surface waters of the FSC may have access to a greater variety of hydrocarbons when grown on dispersed oil (for example in the CEWAF treatment), as observed elsewhere [44], or the *in situ* cold temperatures (∼10°C) and/or nutrient limitation in the FSC may have delayed their response.

One of the early bloomers, *Cycloclasticus*, a microbe known for its appetite for PAHs [12, 37], appeared by day 7 in the WAF treatment predominantly, where it increased in abundance until day 28. *Cycloclasticus* also appeared in BEWAF treatment towards the end of the incubation. *Cycloclasticus* was not observed in the Finasol-amended treatments, which was surprising since previous studies showed *Cycloclasticus* domination of the microbial communities in CEWAF amendments to northeast Atlantic seawater [8, 33] and in natural deepwater oil plumes during the early phase of the Deepwater Horizon spill [39], though which had used different types of a synthetic dispersant. Comparing the dynamics of *Cycloclasticus* across the Finasol- and biosurfactant-amended treatments, it was clear that Finasol negatively impacted this taxon. This inhibition of *Cycloclasticus* has profound implications for oil biodegradation since it is responsible for biodegradation of aromatic hydrocarbons [12]; the absence of *Cycloclasticus* translated to reduced biodegradation rates for the aromatic fraction in CEWAF treatments.

We observed relatively high abundance of an uncultured member of the family *Micavibrionaceae* in the *in situ* FSC microbial community, initially across all treatments, in a later bloom in the BEWAF and SWBS treatments, and to a lesser extent in the WAF (14%) by days 14 and 28. The order *Micavibrionales* is assigned to a group of obligate predatory bacteria, the *Bdellovibrio* and like organisms (BALOs) [45]. Although species from this order were first described in 1982 [46], not much is known about them or their role in natural ecosystems. A recent study from Lake Geneva [47] demonstrated that *Micavibrionaceae* vary throughout the year, with higher numbers in the spring, likely linked to phytoplankton dynamics since they are possible prey for BALOs. Their abundance here may relate to the time of year as our sampling coincided with a phytoplankton bloom. The dynamics of *Micavibrionaceae* may also result from their preying on blooming oil-degrading bacteria, as their abundance increase coincided with decreasing numbers of *Colwellia* and *Oleispira* in the WAF, BEWAF and SWBS treatment by days 14 and 28. Such top-down grazing control of oil degrading organisms is not usually considered when assessing their dynamics, but it is clear that this may be quite important in influencing the composition and biodegradation capability of oil degrading microbial communities.

### Ecological processes drive local community composition

Ecological processes governing the community compositions have been studied for subsurface [48] and desert microbiomes [49] but not for marine microcosms amended with crude oil and surfactants. Overall, we found that mean NTI across all communities was significantly greater than +2 providing evidence that environmental filtering strongly determines local community composition of surface seawater communities when enriched with crude oil in combination with Finasol or rhamnolipid.

Therefore, it seems that the crude oil, dispersant and/or rhamnolipid limit community membership whereby closely related (e.g. belonging to the same family) and ecologically similar (e.g. hydrocarbon-degrading) taxa are more likely to coexist than expected if random ecological processes (drift) assembled the composition. The environmental filtering, however, was strongest in the Finasol-amended communities which also displayed the lowest diversity the incubation period. One explanation is that Finasol dispersed the oil to a greater extent and more of the crude oil compounds are dissolved in the microcosms than the rhamnolipid-amended and oil-only treatments, hence prompting a quicker and stronger response of members of the *Rhodobacteracaea* and *Vibrionaceae* families, known opportunistic oil degraders [12].

### Hydrocarbon degradation

Hydrocarbon analysis revealed faster biodegradation of alkanes in the CEWAF treatment, as indicated by the decreased *n*C_17_/pristane and *n*C_18_/phytane ratios and chromatogram traces. The dominant taxa in the BEWAF and CEWAF treatments were similar on day 7, with the exception for *Cowellia* and *Oleispira* whose abundance in CEWAF was lower than in BEWAF, suggesting that a major portion of the alkane biodegradation had occurred already by day 3 in the CEWAF treatment and was still on-going in the BEWAF treatment. On the last day of the experiment (day 28), the high abundance of *Cycloclasticus* in the WAF and BEWAF treatments suggested on-going PAH biodegradation. In contrast, by day 28 in the CEWAF treatment the community was dominated by members of the *Rhodobacteriaceae* (∼55% of the total community) that may have been involved in utilising the ‘leftovers’ of the earlier alkane-degrading bloom rather than on-going PAH degradation. PAH degradation appears to be limited in the CEWAF treatment, as observed in other studies [4], and this may be due to inhibition of PAH degraders (e.g. *Cycloclasticus*) in this treatment.

The persistent presence of *Alcanivorax* and *Pseudomonas*, albeit at relatively low abundances (4% and 3.7%, respectively), suggests on-going alkane biodegradation through the end of the experiment. The faster biodegradation of alkanes in CEWAF could be explained by the Finasol increasing the bioavailability of these hydrophobic components, more so than the rhamnolipid, and hence more alkanes may have been bioavailable to the hydrocarbonoclastic population when Finasol was used. Our findings support observations of other microbial biodegradation studies where synthetic dispersants enhanced biodegradation of some oil components [8]. A previous study [50] found a substantial decrease in the amount (approx. 3 times) of oil in oil plus dispersant mixtures compared to undispersed oil only, even though biodegradation occurred in both treatments. Similar observations were made in Arctic waters of temperatures down to -1°C, where the indigenous Arctic microbial community could completely degrade *n*-hexadecane, *n*-octadecane and some individual PAHs regardless of the presence of Corexit, but where the biodegradation of the oil (2.5 ppm fresh and weathered), in general, was faster in the presence of the dispersant [7].

## Conclusions

Synthetic dispersants have been used to treat large scale marine oil spills for many years. But whether they enhance microbial hydrocarbon degradation is still under active investigation by the scientific community. Biosurfactants have been proposed as a more environmentally compatible and less toxic alternative to synthetic dispersants. This study aimed to advance the understanding of how rhamnolipid biosurfactant affect oil-contaminated seawater microbial communities compared to synthetic dispersant Finasol, and whether rhamnolipids are suitable for use in oil spill response. Our data support that in the beginning of incubation, the indigenous microbial communities exposed to oil, to oil with dispersant, and to oil with rhamnolipid are similar in composition, all dominated by a diverse set of known oil degrading genera. However, over time, the microbial succession patterns dramatically changed, whereby Finasol suppressed *Cycloclasticus*, but favoured *Vibrio* and *Pseudophaeobacter*. In contrast, the rhamnolipid sustained *Oleispira* and *Micavibrionaceae* abundances. We also found that Finasol stimulated faster *n*-alkane degradation similarly to previous studies, but it suppressed biodegradation of the aromatic fraction which corroborates with the suppression of the obligate aromatic hydrocarbon degrader *Cycloclasticus*. Moreover, the highly deterministic nature of the communities in this study highlights the sensitivity of marine microbes to environmental change.

## Availability of data and materials

The raw sequences files supporting the results of this article are available in the NCBI Sequence Read Archive under accession number PRJNA636672.

## Supporting information

Supplementary Results

## Acknowledgments

We thank Alejandro Gallego from Marine Scotland Science and the crew of MRV *Scotia* for their technical and logistical support to accommodate our research needs during the research cruise to the FSC. We also thank Angelina Angelova for her valuable guidance with the Illumina MiSeq sequencing protocol and analysis, Onoriode Esegbue and Joe Casillo (both Heriot-Watt University) for assistance with GC-FID/MS analysis, and Ibrahim Banat (Ulster University) for providing the rhamnolipid biosurfactant. We would also like to thank Mr John-Philippe Robinson (Total Fluides) for providing the synthetic dispersant Finasol OSR52 and BP for the Schiehallion crude oil.

## Funding

This manuscript contains work conducted during a PhD study undertaken as part of the Natural Environment Research Council (NERC) Centre for Doctoral Training (CDT) in Oil and Gas (NE/M00578X/1). It is sponsored by Heriot-Watt University via their James-Watt Scholarship Scheme to CN and whose support is gratefully acknowledged. Partial support was also provided by the Oil & Gas UK to TG, a NERC Independent Research Fellowship (NERC NE/L011956/1) to UZI, and by an Emmy-Noether fellowship grant (number 326028733) from the German Research Foundation (DFG) awarded to SK.

## Contributions

CN, SK and TG designed this study. CN collected the field samples, performed the experiments and together with CM generated all the data. UZI wrote the analysis script to generate the figures and, together with CN, performed the bioinformatics and statistical analysis. CN, SJ and TG wrote the manuscript, and UZI, CM and SK contributed to its final revision.

## Conflict of Interest

The authors declare no conflict of interest.

## Supplementary Information

**Additional File 1: Supplementary Methods**.**docx**

: Supplementary Methods and Materials used to produce the main results in this study.

**Additional File 2: Supplementary File 2_Top 25 taxa relative abundance**.**xlsx**

: Relative abundance of top 25 taxa in all treatments (including replicates) over time.

**Additional File 3: Supplementary Figure S1**.**pdf**

: Temporal alpha diversity indices of ASVs in treatments over time. Statistically different treatments (pair-wise ANOVA) are connected by bracket and the level of significance is shown with: * (p<0.05), ** (p<0.01), or *** (p<0.001).Treatments are represented by shape (shown on graph) and incubation time by colour: red – baseline microbial community at time of seawater sampling, olive green – day 0, green – day 3, blue – day 7, pink – day 14, and brown – day 28.

**Additional File 4: Supplementary Results**.**docx**

: Additional results to support the results presented in the main article. Includes Fig S2 and Fig S3, and Tables S1-7.

**Additional File 5: Supplementary File 5_Differential abundant taxa**.**xlsx**

: DESeq2 results for the differential expressed bacterial taxa between treatments BEWAF, WAF and CEWAF.

**Additional File 6: Supplementary Figure S4**.**pdf**

: Differential heat trees showing the key (significant) differential taxa (DESeq2; Wilcoxon p-value test adjusted with multiple comparison) in seawater-only control treatment (SW). The top 3 subsets with the highest correlation with the full ASV table considering Bray-Curtis distance (PERMANOVA) are listed for each treatment. The grey tree is the taxonomy key for the smaller unlabelled coloured trees. The colour of each taxon represents the log-10 ratio of median proportions of reads observed in each treatment. The size of tree nodes shows the number of ASVs (here labelled as OTUs) present in each treatment.

**Additional File 7: Supplementary Figure S5**.**pdf**

: Temporal measures for nearest-relative index (NRI) and nearest-taxon index (NTI. Colours represent incubation time: red – baseline microbial community at time of seawater sampling, olive green – day 0, green – day 3, blue – day 7, pink – day 14, and brown – day 28. Statistically different treatments (pair-wise ANOVA) are connected by bracket and the level of significance is shown with: * (p<0.05), ** (p<0.01), or *** (p<0.001).

**Additional File 8: Supplementary Figure S6**.**pdf**

: Overall predicted functional alpha diversity of microbial pathways (expressed as number of KEGG orthologs). Statistically different treatments (pair-wise ANOVA) are connected by bracket and the level of significance is shown with: * (p<0.05), ** (p<0.01), or *** (p<0.001). Colours represent treatments and shapes the incubation time.

**Additional File 9: Supplementary Figure S7**.**pdf**

: (**A)** Predicted functional alpha diversity of microbial pathways (expressed as number of KEGG orthologs). Statistically different treatments (pair-wise ANOVA) are connected by bracket and the level of significance is shown with: *(p<0.05), ** (p<0.01), or *** (p<0.001). **(B)** Principal coordinate analysis (PCoA) on beta diversity measured with Bray-Curtis dissimilarity distance matrix. In both (A) and (B) treatments are represented by shape (shown on graph) and incubation time by colour: red – baseline microbial community at time of seawater sampling, olive green – day 0, green – day 3, blue – day 7, pink – day 14, and brown – day 28.

**Additional File 10: Supplementary Figure S8**.**pdf**

: Heatmap showing the scaled log abundance (color key on top left) of aliphatic and aromatic degradation, and biosurfactant synthesis pathways. Pathways are shown along the y-axis and BEWAF, CEWAF, and WAF samples along the x-axis. The two color-coded bars on top of the heatmap indicate their treatments and incubation days status. Hierarchical clustering of the samples (top) is based on the correlation between samples’ predicted gene expression.

**Additional File 11: Supplementary Figure S9**.**pdf**

: Differences in aliphatic and polycyclic aromatic hydrocarbon biomarker ratios of three different treatments over time in days (grey boxes): nC17/pristane (nC17/pri), nC18/phytane (nC18/phy), Phenanthrene/9-methylphenanthrene (P/9MP), (3+2)-methylphenanthrene/(9+1)-mehylphenanthrene (3+2MP/9+1MP), and 3-methylphenanthrene/9-methylphenanthrene (3MP/9MP). Values are the mean of three independent replicates (except for BEWAF day 0 (two replicates) and CEWAF day 28 (one replicate)) +/- standard deviation.

## Notes

### Competing Interest Statement

The authors have declared no competing interest.

## References

1. Joye S, Kostka J. Microbial genomics of the global ocean system. Earth and Space Science Open Archive. 2020.

2. National Academies of Sciences, Engineering and M. The Use of Dispersants in Marine Oil Spill Response. Washington, D.C.: National Academies Press; 2020.

3. Hamdan LJ, Fulmer PA. Effects of COREXIT EC9500A on bacteria from a beach oiled by the Deepwater Horizon spill. Aquatic Microbial Ecology. 2011;63:101–9.

4. Kleindienst S, Seidel M, Ziervogel K, Grim S, Loftis K, Harrison S, et al. Chemical dispersants can suppress the activity of natural oil-degrading microorganisms. Proceedings of the National Academy of Sciences. 2015;112:14900–5.

5. Rahsepar S, Smit MPJ, Murk AJ, Rijnaarts HHM, Langenhoff AAM. Chemical dispersants: Oil biodegradation friend or foe? Marine Pollution Bulletin. 2016;108:113–9.

6. Prince RC, McFarlin KM, Butler JD, Febbo EJ, Wang FCY, Nedwed TJ. The primary biodegradation of dispersed crude oil in the sea. Chemosphere. 2013;90:521–6.

7. McFarlin KM, Prince RC, Perkins R, Leigh MB. Biodegradation of dispersed oil in Arctic seawater at -1°C. PLoS ONE. 2014;9:1–8.

8. Brakstad OG, Ribicic D, Winkler A, Netzer R. Biodegradation of dispersed oil in seawater is not inhibited by a commercial oil spill dispersant. Marine Pollution Bulletin. 2018;129:555–61.

9. Judson RS, Martin MT, Reif DM, Houck KA, Knudsen TB, Rotroff DM, et al. Analysis of eight oil spill dispersants using rapid, in vitro tests for endocrine and other biological activity. Environmental Science and Technology. 2010;44:5979–85.

10. Kujawinski EB, Soule MCK, Valentine DL, Boysen AK, Longnecker K, Redmond MC. Fate of Dispersants Associated with the Deepwater Horizon Oil Spill. Environmental Science and Technology. 2011;45:1298–306.

11. Campo P, Venosa AD, Suidan MT. Biodegradability of Corexit 5900 and Dispersed South Louisiana Crude Oil at 5C and 25C. Environmental Science and Technology. 2013;47:1960–7.

12. Head IM, Jones DM, WFM Röling. Marine microorganisms make a meal of oil. Nature Reviews Microbiology. 2006;4:173–82.

13. Das P, Yang XP, Ma LZ. Analysis of biosurfactants from industrially viable Pseudomonas strain isolated from crude oil suggests how rhamnolipids congeners affect emulsification property and antimicrobial activity. Frontiers in Microbiology. 2014;5 DEC:1–8.

14. Nikolopoulou M, Eickenbusch P, Pasadakis N, Venieri D, Kalogerakis N. Microcosm evaluation of autochthonous bioaugmentation to combat marine oil spills. New Biotechnology. 2013;30:734– 42.

15. Chen Q, Bao M, Fan X, Liang S, Sun P. Rhamnolipids enhance marine oil spill bioremediation in laboratory system. Marine Pollution Bulletin. 2013;71:269–75.

16. Couto CR de A, Jurelevicius D de A, Alvarez VM, van Elsas JD, Seldin L. Response of the bacterial community in oil-contaminated marine water to the addition of chemical and biological dispersants. Journal of Environmental Management. 2016;184:473–9.

17. Gallego A, O’Hara Murray R, Berx B, Turrell WR, Beegle-Krause CJ, Inall M, et al. Current status of deepwater oil spill modelling in the Faroe-Shetland Channel, Northeast Atlantic, and future challenges. Marine Pollution Bulletin. 2018;127 January:484–504.

18. Bett BJ. UK atlantic margin environmental survey: Introduction and overview of bathyal benthic ecology. Continental Shelf Research. 2001;21:917–56.

19. Tillett D, Neilan BA. Xanthogenate nucleic acid isolation from cultured and environmental cyanobacteria. Journal of Phycology. 2000;36:251–8.

20. Berry D, Ben Mahfoudh K, Wagner M, Loy A. Barcoded primers used in multiplex amplicon pyrosequencing bias amplification. Applied and environmental microbiology. 2011;77:7846–9.

21. Bolyen E, Rideout JR, Dillon MR, Bokulich NA, Abnet CC, Al-Ghalith GA, et al. Reproducible, interactive, scalable and extensible microbiome data science using QIIME 2. Nature Biotechnology. 2019;37:852–7.

22. Callahan BJ, McMurdie PJ, Rosen MJ, Han AW, Johnson AJA, Holmes SP. DADA2: High-resolution sample inference from Illumina amplicon data. Nature Methods. 2016;13:581–3.

23. Callahan BJ, McMurdie PJ, Holmes SP. Exact sequence variants should replace operational taxonomic units in marker-gene data analysis. ISME Journal. 2017;11:2639–43.

24. Quast C, Pruesse E, Yilmaz P, Gerken J, Schweer T, Yarza P, et al. The SILVA ribosomal RNA gene database project: Improved data processing and web-based tools. Nucleic Acids Research. 2013;41:590–6.

25. Douglas GM, Maffei VJ, Zaneveld J, Yurgel SN, Brown JR, Taylor CM, et al. PICRUSt2: An improved and extensible approach for metagenome inference. bioRxiv. 2019;:672295.

26. R Core Team. R: A language and environment for statistical computing. 2019.

27. Marchant R, Banat IM. Microbial biosurfactants: Challenges and opportunities for future exploitation. Trends in Biotechnology. 2012;30:558–65.

28. Ron EZ, Rosenberg E. Enhanced bioremediation of oil spills in the sea. Current Opinion in Biotechnology. 2014;27:191–4.

29. Yakimov MM, Giuliano L, Gentile G, Crisafi E, Chernikova TN, Abraham WR, et al. Oleispira antarctica gen. nov., sp. nov., a novel hydrocarbonoclastic marine bacterium isolated from Antarctic coastal sea water. International Journal of Systematic and Evolutionary Microbiology. 2003;53:779– 85.

30. Yakimov MM, Timmis KN, Golyshin PN. Obligate oil-degrading marine bacteria. Current Opinion in Biotechnology. 2007;18:257–66.

31. Ribicic D, Netzer R, Winkler A, Brakstad OG. Microbial communities in seawater from an Arctic and a temperate Norwegian fjord and their potentials for biodegradation of chemically dispersed oil at low seawater temperatures. Marine Pollution Bulletin. 2018;129:308–17.

32. Kleindienst S, Grim S, Sogin M, Bracco A, Crespo-Medina M, Joye SB. Diverse, rare microbial taxa responded to the Deepwater Horizon deep-sea hydrocarbon plume. The ISME Journal. 2016;10:1–16.

33. Suja LD, Summers S, Gutierrez T. Role of EPS, Dispersant and Nutrients on the Microbial Response and MOS Formation in the Subarctic Northeast Atlantic. Frontiers in Microbiology. 2017;8 April:1–15.

34. Gontikaki E, Potts LD, Anderson JA, Witte U. Hydrocarbon-degrading bacteria in deep-water subarctic sediments (Faroe-Shetland Channel). Journal of Applied Microbiology. 2018;125:1040–53.

35. Hazen TC, Dubinsky EA, DeSantis TZ, Andersen GL, Piceno YM, Singh N, et al. Deep-Sea Oil Plume Enriches Indigenous Oil-Degrading Bacteria. Science. 2010;330:204–8.

36. Valentine DL, Kessler JD, Redmond MC, Mendes SD, Heintz MB, Farwell C, et al. Propane respiration jump-starts microbial response to a deep oil spill. Science. 2010;330:208–11.

37. Gutierrez T, Singleton DR, Berry D, Yang T, Aitken MD, Teske A. Hydrocarbon-degrading bacteria enriched by the Deepwater Horizon oil spill identified by cultivation and DNA-SIP. The ISME journal. 2013;7:2091–104.

38. Hazen TC, Prince RC, Mahmoudi N. Marine Oil Biodegradation. Environmental Science and Technology. 2016;50:2121–9.

39. Hu P, Dubinsky EA, Probst AJ, Wang J, Sieber CMK, Tom LM, et al. Simulation of Deepwater Horizon oil plume reveals substrate specialization within a complex community of hydrocarbon degraders. Proceedings of the National Academy of Sciences of the United States of America. 2017;114:7432–7.

40. Abisado RG, Benomar S, Klaus JR, Dandekar AA, Chandler JR. Bacterial quorum sensing and microbial community interactions. mBio. 2018;9:1–13.

41. Chakraborty R, Borglin SE, Dubinsky EA, Andersen GL, Hazen TC. Microbial response to the MC-252 oil and Corexit 9500 in the Gulf of Mexico. Frontiers in Microbiology. 2012;3 OCT:1–6.

42. Techtmann SM, Zhuang M, Campo P, Holder E, Elk M, Hazen TC, et al. Corexit 9500 enhances oil biodegradation and changes active bacterial community structure of oilenriched microcosms. Applied and Environmental Microbiology. 2017;83:1–14.

43. Watson JS, Jones DM, Swannell RPJ, ACT Van Duin. Formation of carboxylic acids during aerobic biodegradation of crude oil and evidence of microbial oxidation of hopanes. Organic Geochemistry. 2002;33:1153–69.

44. Overholt WA, Marks KP, Romero IC, Hollander DJ, Snell TW, Kostka JE. Hydrocarbon-degrading bacteria exhibit a species-specific response to dispersed oil while moderating ecotoxicity. Applied and Environmental Microbiology. 2016;82:518–27.

45. Davidov Y, Huchon D, Koval SF, Jurkevitch E. A new α-proteobacterial clade of Bdellovibrio-like predators: Implications for the mitochondrial endosymbiotic theory. Environmental Microbiology. 2006;8:2179–88.

46. Lambina, V. A, Afinogenova A V., Penabad SR, Konovalova SM, Pushkareva AP. Micavibrio admirandus gen. et sp. nov. Mikrobiologiya. 1982;51:114–7.

47. Ezzedine JA, Jacas L, Desdevises Y, Jacquet S. Bdellovibrio and Like Organisms in Lake Geneva: An Unseen Elephant in the Room? Frontiers in Microbiology. 2020;11:1–14.

48. Stegen JC, Lin X, Konopka AE, Fredrickson JK. Stochastic and deterministic assembly processes in subsurface microbial communities. ISME Journal. 2012;6:1653–64.

49. Caruso T, Chan Y, Lacap DC, Lau MCY, McKay CP, Pointing SB. Stochastic and deterministic processes interact in the assembly of desert microbial communities on a global scale. ISME Journal. 2011;5:1406–13.

50. Prince RC, Butler JD. A protocol for assessing the effectiveness of oil spill dispersants in stimulating the biodegradation of oil. Environmental Science and Pollution Research. 2014;21:9506– 10.

