## Supplementary Methods for "Response and oil degradation activities of a northeast Atlantic bacterial community to biogenic and synthetic surfactants"

### Field sample collection in the Faroe-Shetland Channel

During a research cruise on MRV *Scotia* in May of 2018, surface seawater (3m depth) was collected from the Faroe-Shetland Channel (FSC), a subarctic region of the northeast Atlantic. The seawater temperature at the time of collection was measured at 9.7 °C and the salinity 35.28‰. This sampling site lies on the Fair Isle-Munken line [1] near the Foinaven oil field development area, approximately 3 and 9.3 nautical miles respectively from the Petrojarl Foinaven and Glen Lion production facilities. Collection of seawater samples was performed using 10 L Niskin water bottles mounted on a CTD (conductivity, temperature, depth) carousel. Immediately after recovery, some of the collected seawater was used to rinse, at least three times, two Nalgene carboys (10L each; acid-washed, acetone-rinsed and dried) prior to filling, and immediately stored at 10 °C onboard the vessel for transport back to the laboratory for preparation of water-accommodated fractions (WAFs) and for the crude oil and biosurfactant/dispersant enrichment experiments, as described below. Sub-samples were collected for DNA extraction and quantification of the *in-situ* bacterial communities and to account for any changes before initiation of experiments (described below). For this, 3 litres of the sampled water were filtered through polycarbonate membrane filters (1L per filter; 0.22μm pore size) and the filters stored onboard at -20 °C until return to the laboratory when they were transferred to -80 °C for subsequent analysis.

### Water accommodated fractions

Three main water accommodated fractions (WAFs) were prepared in acetone-rinsed, acid-washed and autoclaved 2 L glass aspirator bottles (with an outlet at the bottom) according to previous methods [2, 3], though with some modifications as described below. For preparation of the WAFs, the seawater was filtered (0.22μm; Millipore) in order to avoid the possibility of bacterial growth during the preparation of the WAFs. The first WAF contained seawater and crude oil only and is on referred to as WAF.  A Chemically Enhanced WAF (CEWAF) was prepared with seawater, crude oil and addition of the synthetic dispersant Finasol OSR-52 (Total Fluides, Paris, France) at a dispersant-to-oil (DOR) ratio of 1:20. Biosurfactant Enhanced WAF (BEWAF) was prepared with seawater, crude oil and rhamnolipid (produced by *P. aeruginosa*) at the same DOR as in the CEWAF. All three WAFs contained the same volume of filter-sterilised seawater (1560 ml) and Schiehallion crude oil (120 ml; API 25°; BP) which also originates from the FSC. The working concentration of the rhamnolipid used to prepare the BEWAF was 10 000 mg/L with critical micelle concentration of 27 mN/m. Briefly, each of the three main WAFs (WAF, CEWAF, BEWAF) were prepared by combining the prescribed quantities of seawater, crude oil and synthetic dispersant or biosurfactant in the aspirator bottles and leaving the solutions to mix on a rotary magnetic stirrer (140 rpm; 10°C) for up to 48 hours. In addition, two control WAFs were set up in the same way to assess the microbial community response to the dispersant or biosurfactant alone and in the absence of the crude oil. One of these contained only seawater and Finasol (SWD); the other contained seawater and rhamnolipid (SWBS). Both Finasol and rhamnolipid were used at the same concentration as in the CEWAF and BEWAF, respectively. After mixing for 48 hours, the three WAFs containing crude oil were allowed to stand undisturbed for one hour to allow for any undispersed oil to settle to the surface. The aqueous phase from each of the WAF mixtures was then carefully collected from the bottom outlet of the bottles, avoiding any of the undispersed oil.

**Setup and sampling of microcosm treatments**

Acetone-rinsed, acid-washed and autoclaved 500 ml glass bottles, with Teflon-lined caps, were used and each treatment was replicated in triplicate. Each treatment contained 66 ml of filtered WAF, CEWAF, BEWAF, SWD, or SWBS added to 234 ml of unfiltered seawater to a total volume of 300 ml, leaving 200 ml of head space to ensure aerobic conditions. In addition, untreated control comprising seawater alone with no other additions was setup and run in parallel. All of the bottle treatments were placed on a roller table to maintain constant gentle mixing (15 rpm) and at temperature of 9.7 °C (*in situ* temperature at the time of sampling) for 4 weeks in darkness. At the beginning of these incubations (day 0), and then subsequently thereafter at days 3, 7, 14 and 28, each treatment was sub-sampled (5 ml) for total microbial cell counts, and also for DNA extraction (10 ml), following the methods described below. Additional WAF, BEWAF and CEWAF microcosms (in triplicate) were set up identically to the rest of the microcosms for hydrocarbon biodegradation analysis as described below.

**DNA extraction and barcoded amplicon 16S rRNA gene sequencing**

Filters with collected biomass from each treatment (incl. replicates) were each transferred into sterile 1.5 ml Eppendorf tubes and submerged in liquid nitrogen until completely frozen. The frozen filters were carefully crushed into fine particles using sterile pipette tips. DNA was extracted according to the method of a previous study [4]. Sterile deionised water samples were included to act as negative controls to qualify the sterility of the method and of the reagents used. DNA extracts were resuspended in 20 µl of 1 mM TE buffer and confirmed by gel electrophoresis prior to storage at -20 °C for Illumina barcoded-amplicon sequencing.

A two-step amplification procedure was used to minimize the heteroduplex formation in mixed-template reactions [5]. The 16S rRNA gene was first amplified with universal bacterial primers 8F (5`-AGAGTTTGATCCTGGCTCAG-3`) [6] and 1492R (5`-TACGGYTACCTTGTTACGACT-3`) [7] in triplicate 25 µl reactions each containing 2.5 U MyTaq™ Red DNA Polymerase (Bioline Reagents Ltd.), 5 µl 5x MyTaq Red Reaction buffer (containing 5 mM dNTPs and 15 mM $MgCl_2$) (Bioline Reagents Ltd.), 17.5 µl molecular grade water, forward and reverse primer at final concentration of 0.2 µM each, and 1 µl of target DNA. Amplification was carried out on a thermocycler using the following conditions: initial denaturation at 96 °C for 5 min, followed by 32 cycles of 96 °C for 30 sec, 54 °C for 30 sec and 72 °C for 30 sec, and a final extension at 72 °C for 10 min. Purified samples were quantitated with Nanodrop and stored at -20 °C for the second step of barcoded amplification.

The second step involved the targeted barcoded amplification of the V4 hypervariable region of the 16S rRNA gene from the first-step PCR. For each sample, this was performed using duplicate 25 µl reactions. Each reaction consisted of the same reagents as in the first PCR step but this time barcoded 515F (5′-GTGYCAGCMGCCGCGGTAA-3′) [8] and 806R (5′-GGACTACNVGGGTWTCTAAT-3′) [9] primers were added to the PCR mixture instead of 8F and 1492R primers. Both primers contained Illumina MiSeq adapters added to the 5′ ends and unique Golay barcodes added to the 515F primer. Barcoded amplification was carried out as per Earth Microbiome project's thermocycler conditions [10].

**Bioinformatics and statistical analyses**

Alpha diversity indices Shannon and Species Richness were calculated on the rarefied-to minimum-reads ASVs to investigate within-community diversity of each treatment through time. Variance between treatments at different incubation times for each alpha diversity index were compared by performing ANOVA. Community composition variation between treatments and incubation time (Beta diversity) were assessed using pairwise Permutational Multivariate Analysis of Variance (PERMANOVA) and the adonis function in vegan package [11]. Principal Coordinate Analysis plots (PCoA) were used to visualise dissimilarities based on Bray-Curtis similarity matrix (considers the species abundance count), Unweighted UniFrac (considers phylogenetic distance between branch lengths of ASVs observed in different samples without taking into account the abundances) and Weighted UniFrac distance (unweighted unifrac distance weighted by the abundances of ASVs) matrices calculated from the rarefied and normalised (relative abundance) ASV table. Significance of results was based on high F values and level of significance based on low p-values (<0.05) for a comparison to be considered a significantly different.

To understand multivariate homogeneity of groups, variances between multiple conditions (combinations of treatments over time), vegan's betadisper function was used, in which the distances between objects and group centroids are handled by using a reduced order representation based on distance metrics (Bray-Curtis, Unweighted UniFrac, or Weighted UniFrac) in principal coordinates space and afterwards performing ANOVA on differences of each sample from the mean of the group they belong to. Vegan's adonis function was used for multivariate analysis of variance (PERMANOVA) among sources of variation (treatment and sampling time) using distance matrices (Bray-Curtis/Unweighted Unifrac/Weighted Unifrac). To determine which ASVs were significantly different between multiple conditions (treatments/days), DESeq2 package [12] was used. This function uses negative binomial GLM-fitting and wald statistics prior to applying Bayesian shrinkage.

To show how each sample was markedly different in terms of beta diversity, we have performed Local Contribution to Beta Diversity (LCBD) analysis using adespatial package [13]. The procedure calculates the total beta diversity considering all the samples, and then allocates the proportions based on how different the microbial community structure of a single sample is from the average (with higher LCBD values representing outliers) beta diversity, and also provides a mean to show when the community structure has stabilised in a temporal setting [14]. To characterize the phylogenetic community composition within each sample, whether the microbial community structure is driven by competition among taxa (without any extrinsic environmental impact) or being deterministic, we quantified the nearest-taxon-index (NTI; local phylogenetic clustering) and nearest-relative-index (NRI; global phylogenetic clustering).

We adopted the approach by Stegen et al. [15] and used recently in a chicken microbiom study [14].

We performed subset regression of different metrics of microbiome (alpha and beta diversity) by testing all possible combination of the predictor variables (in our case, categorical variables), and then selecting the best model according to some statistical criteria, with recommendations given by Kassambra et al. [16] with their code available online here. The R function regsubsets from leaps package [17] was used to identify different best models of different sizes, by specifying the option nvmax, set to the maximum number of predictors to incorporate in the model. Having obtained the best possible subsets, the k-fold cross-validation consisting of first dividing the data into k subsets. Each subset (10%) served successively as test data set and the remaining subset (90%) as training data. The average cross-validation error was then computed as the model prediction error. This was computed using a custom function utilising R's train function from the caret package [18] and tab_model function from sjPlot package [19].

To categorise the predicted functional diversity, we used KEGG (Kyoto Encyclopedia of Genes and Genomes) pathway mapping tool and MetaCyc databases [20]. KEGG orthologs (KO) involved in aliphatic and aromatic hydrocarbon degradation were manually filtered along with biosurfactant-associated enzymes for each treatment. We specifically looked at genes for rhamnolipid (glycolipid) and surfactin (lipopeptide) synthesis as they are the most studied and thesse metabolic pathways well known. Because our study focused on a marine microbial community, we also selected genes for exopolymer production which are more characteristic in marine bacteria, for example *Alcanivorax* sp. and *Halomonas* sp., and also genes encoding the *rhl* quorum sensing system in rhamnolipid-producing strains that regulates the production of rhamnolipid biosurfactants.

**Hydrocarbon degradation analysis by GC-FID/MS**

Replicate bottles (300-ml volume per bottle) were sacrificed for hydrocarbon extraction at three time points – on days 0, 7 and 28. Total hydrocarbons were extracted via solvent extraction using dichloromethane (DCM; HPLC grade, ThermoFisher, UK) at an oil/water mix to DCM ratio of 1:1. The DCM fraction was removed into acid-washed and dried pre-weighted 500-ml round-bottom flasks and the aqueous phase re-extracted with DCM twice more.

Combined DCM extracts were then rotary evaporated to ~2 ml at 40°C and transferred to amber glass vials and stored at -20 °C for subsequent analysis by GC-FID/MS.

A known aliquot corresponding to ca. 10 mg of total hydrocarbon content was taken from each sample and transferred to a 10 ml vial and dried down using a gentle stream of nitrogen gas. Sample residues were dissolved in a small volume of hexane (~200 µl). A methodological blank (no crude oil) was used to ensure no contamination from any residual hydrocarbons during the analysis, and a sample of the original Schiehallion crude oil was analysed as a reference. Total hydrocarbon extracts were separated into two complementary polarity-defined fractions (aliphatic and aromatic) by using so-called *flash* open-column chromatography over 0.5 g silica gel (0.060 – 0.200mm, 60A) and 0.5 g alumina (aluminium oxide, 50-200µm, 60A) sorbents. Sorbents were pre-extracted with DCM to minimize organic contaminants and then activated at 120 °C prior to use. Sorbents were introduced to open columns as slurries in hexane. Each column was packed with half the alumina, topped with silica gel, and then alumina again. The column was flushed with at least two bed-volumes of hexane (~4 ml) before the sample, which was dissolved in hexane, was applied to the top of the column. The aliphatic fraction of the total petroleum hydrocarbons (TPH) was eluted with 4 ml of hexane into an organics-free 2 ml glass vial. The aromatic fraction of the TPH was eluted with 4 ml DCM into a separate vial. Elution solvents were dried-down under nitrogen, re-dissolved in hexane (aliphatic fraction) and DCM (aromatic fraction) and transferred to organics-free 300 µl GC/MS borosilicate vial inserts. The samples were dried as before with nitrogen gas. The TPH fractions were then analysed by injecting 1 µl of the hydrocarbon fraction diluted in hexane in the autosampler of a Thermo Trace 1310 GC coupled with Thermo ISQ LT MS and fitted with a splitless constant temperature injector (300 °C), a flame ionisation detector (FID) at 310 °C, and an HP-5MS capillary column (30m x 0.25mm x 0.2µm; Agilent). The column programme was set at 50 °C for 2 min and 5 °C/min to 310 °C for 21 min to a total run time of 75 minutes. Chromatographic data were acquired and processed through Chromeleon software (v. 7.2.8; Thermo Fisher). Peak areas of individual $C_{12}$ to $C_{30}$ *n*-alkanes and the isoprenoids pristane and phytane were calculated. The aromatic hydrocarbons were analysed by GC-MS in full scan mode (50-600 amu at 4 min).

Peak areas of aliphatic and aromatic hydrocarbon species/groups that were biodegraded after 7 or 28 days were calculated by subtracting the respective hydrocarbon concentrations measured in the control from those of the treatment incubations. Additionally, ratios of *n*-alkanes to acyclic isoprenoid hydrocarbons ($nC_{17}$/pristane and $nC_{18}$/phytane) were used as conventional indicators of biological degradation, due to the recalcitrance imparted by the branched structure of the isoprenoid biomarkers [21]. Similarly, for the aromatic hydrocarbons, four ratios indicative of biodegradation were determined (phenanthrene/9-methylphenanthrene, 3+2-methylphenanthrene/9+1-methylphenanthrene, 3-methylphenanthrene/9-methylphenanthrene) [21]. Two-way ANOVA and *post-hoc* Tukey tests were performed to test for significant differences in the degradation of the hydrocarbons analysed between the treatments.

### References


1. Turrell WR, Slesser G, Adams RD, Payne R, Gillibrand PA. Decadal variability in the composition of Faroe Shetland Channel bottom water. Deep-Sea Research Part I: Oceanographic Research Papers. 1999;46:1–25.

2. Aurand D, Coelho GM. Cooperative Aquatic Toxicity Testing of Dispersed Oil and the "Chemical Response to Oil Spills: Ecological Effects Research Forum (CROSERF)". Lusby, MD; 2005.

3. Kleindienst S, Seidel M, Ziervogel K, Grim S, Loftis K, Harrison S, et al. Chemical dispersants can suppress the activity of natural oil-degrading microorganisms. Proceedings of the National Academy of Sciences. 2015;112:14900–5.

4. Tillett D, Neilan BA. Xanthogenate nucleic acid isolation from cultured and environmental cyanobacteria. Journal of Phycology. 2000;36:251–8.

5. Berry D, Ben Mahfoudh K, Wagner M, Loy A. Barcoded primers used in multiplex amplicon pyrosequencing bias amplification. Applied and environmental microbiology. 2011;77:7846–9.

6. Galkiewicz JP, Kellogg CA. Cross-Kingdom Amplification Using Bacteria-Specific Primers: Complications for Studies of Coral Microbial Ecology. APPLIED AND ENVIRONMENTAL MICROBIOLOGY. 2008;74:7828–31.

7. Chirino B, Strahsburger E, Agulló L, González M, Seeger M. Genomic and Functional Analyses of the 2-Aminophenol Catabolic Pathway and Partial Conversion of Its Substrate into Picolinic Acid in Burkholderia xenovorans LB400. PLoS ONE. 2013;8.

8. Parada AE, Needham DM, Fuhrman JA. Every base matters: Assessing small subunit rRNA primers for marine microbiomes with mock communities, time series and global field samples.

Environmental Microbiology. 2016;18:1403–14.

9. Apprill A, Mcnally S, Parsons R, Weber L. Minor revision to V4 region SSU rRNA 806R gene primer greatly increases detection of SAR11 bacterioplankton. Aquatic Microbial Ecology. 2015;75:129–37.

10. Caporaso JG, Lauber CL, Walters WA, Berg-Lyons D, Lozupone CA, Turnbaugh PJ, et al. Global patterns of 16S rRNA diversity at a depth of millions of sequences per sample. PNAS. 2011;108 Supplement_1:4516–22.

11. Oksanen AJ, Blanchet FG, Friendly M, Kindt R, Legendre P, Mcglinn D, et al. Community ecology package ‘ vegan ’. R package version 2.5-6. 2019.

12. Love MI, Huber W, Anders S. Moderated estimation of fold change and dispersion for RNA-seq data with DESeq2. Genome Biology. 2014;15:1–21.

13. Dray AS, Blanchet G, Borcard D, Guenard G, Jombart T, Larocque G, et al. Package ‘adespatial’. R package version 0.0-8. 2017.

14. Ijaz UZ, Sivaloganathan L, McKenna A, Richmond A, Kelly C, Linton M, et al. Comprehensive longitudinal microbiome analysis of the chicken cecum reveals a shift from competitive to environmental drivers and a window of opportunity for Campylobacter. Frontiers in Microbiology. 2018;9 OCT:1–14.

15. Stegen JC, Lin X, Konopka AE, Fredrickson JK. Stochastic and deterministic assembly processes in subsurface microbial communities. ISME Journal. 2012;6:1653–64.

16. Kassambara A. Machine Learning Essentials: Practical Guide in R. STHDA; 2018.

17. Lumley T. Leaps: Regression subset selection. R package version 3.1. 2020.

18. Kuhn M. caret Package. Journal Of Statistical Software. 2008;28:1–26.

19. Lüdecke D. sjPlot - Data visualization for statistics in social science. 2014;:2016.

20. Caspi R, Billington R, Keseler IM, Kothari A, Krummenacker M, Midford PE, et al. The MetaCyc database of metabolic pathways and enzymes-a 2019 update. Nucleic Acids Research. 2019;48:445–53.

21. Dawson KS, Schaperdoth I, Freeman KH, Macalady JL. Anaerobic biodegradation of the isoprenoid biomarkers pristane and phytane. Organic Geochemistry. 2013;65:118–26.
