## Supplementary Results for "Response and oil degradation activities of a northeast Atlantic bacterial community to biogenic and synthetic surfactants"

### Supplementary Results and Discussion

#### Results

##### Bacterial diversity variations

Similar to beta diversity analysis, the Local Contribution to Beta Diversity (LCBD) showed each sample's contribution to the observed changes derived as a proportion of the total beta diversity. The diversity patterns with high LCBD values suggested that those samples have markedly different microbial communities. Based on diversity counts (Bray-Curtis, Fig. S2), diversity variation generally increased over time across treatments. At day 0, all treatments show significantly lower, but similar diversity compared to the *in situ* FSC community. As time progressed, the within treatment diversity began to vary and at the end of the incubation period, the CEWAF and SWD treatments displayed the highest contribution to beta diversity variation, and hence their bacterial communities were significantly different from other treatments. Based on unweighted UniFrac, the community of the SWD treatment was significantly distinct from the other treatments, especially on day 7, which coincided with the emergent dominance of members within the family *Rhodobacteraceae*, such as of *Pseudophaeobacter*, *Sedimentitalea* and *Shimia*. In contrast, BEWAF and WAF treatments were dominated by members of the *Colwelliaceae* and *Saccharospirillaceae*.

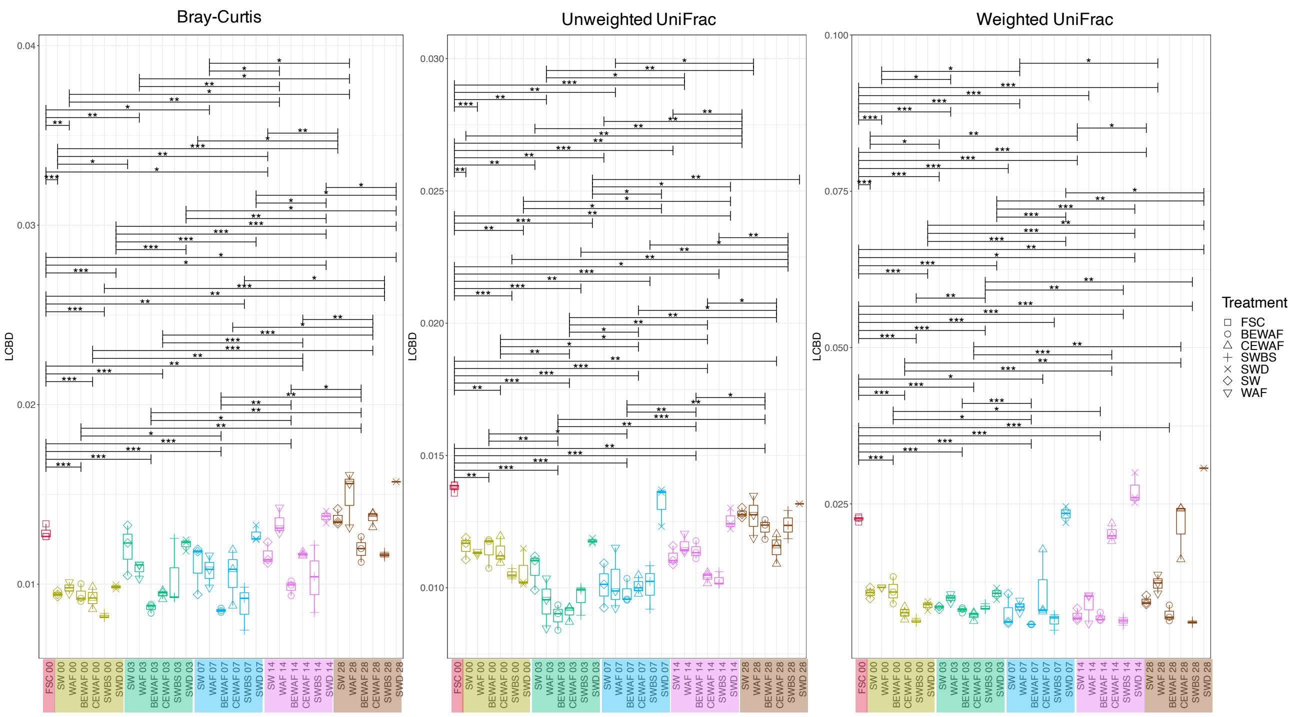


**Figure S2.** Local contribution to beta diversity (LCBD) for Bray-Curits, Unweighted and Weighted UniFrac distance matrices. Colours represent incubation time: red – in situ FSC at time of sampling, olive green – day 0, green – day 3, blue – day 7, pink – day 14 and brown – day 28. Statistically different treatments (pair-wise ANOVA) are annotated with bracket and the level of significance is shown above the bracket with a star sign: * is p<0.05, ** is p<0.01, and *** is p<0.001.

Regression analysis through supervised machine learning showed which variable best explained the variation of diversity in the dataset (Fig. S3). The variables that most significantly affected the alpha and beta diversity variation in the optimal model (with the lowest prediction error) are highlighted in red and blue. The top 10 predictive models for each metric of microbiome analysis – alpha diversity indices (Richness and Shannon), beta diversity (local contribution to beta diversity based on Bray-Curtis, Unweighted and Weighted UniFrac distance matrices), and NTI and NRI, were compiled into tables (Tables S1-7). The variables that positively affected the species richness and Shannon index was in situ FSC and incubation day 0, while incubation days 3 and 7, and treatments CEWAF and SWD, negatively affected alpha diversity variation. The treatments CEWAF, SWD, WAF and BEWAF, as well as incubation day 3, positively affected NRI variation, indicating that these treatments increased clustering of the bacterial communities. Conversely, no clustering was expected on incubation day 0. The presence of Finasol in the seawater (SWD treatment) positively influenced (or increased) the variation in beta diversity (LCBD) of the bacterial community. In terms of alpha diversity, Finasol-amended treatments (CEWAF and SWD) reduced the species richness of the bacterial community. This coincided with the dominance of predominantly four taxa, *Colwellia*, *Oleispira*, *Sedimentitalea* and *Vibrio* (Figure 2 in main text) during the same period. In contrast, the *in situ* FSC community, which was not amended with crude oil, Finasol or the biosurfactant was a positively influencing parameter on alpha and beta diversity. The addition of crude oil alone to seawater (WAF) significantly influenced the variation of diversity in the dataset (LCBD based on Bray-Curtis) as well as the environmental filtering (NTI).

| | Richness | Shannon | NRI | NTI | LCBD Bray-Curtis | LCBD Unweighted UniFrac | LCBD Weighted UniFrac |
|---|---|---|---|---|---|---|---|
| *Treatment_CEWAF* | *** | | *** | | *** | | *** |
| *Treatment_SWD* | *** | ** | *** | * | *** | *** | *** |
| *Treatment_BEWAF* | | ** | *** | | | | |
| *Treatment_SWBS* | | | | | | | |
| *Treatment_SW* | | | | * | *** | ** | |
| *Treatment_WAF* | | | * | | *** | | |
| *Treatment_FSC* | *** | | | ** | *** | *** | *** |
| *Incubation_day_0* | *** | *** | *** | | *** | | |
| *Incubation_day_3* | *** | ** | ** | | | *** | |
| *Incubation_day_7* | *** | | | ** | | | |
| *Incubation_day_14* | | | | | *** | | ** |
| *Incubation_day_28* | | | | | *** | *** | *** |

**Figure S3.** Heatmap of significant extraneous parameters that influence different attributes of microbiome determined by regression analysis. The parameters shown here are from the optimal model for each metric, where blue represent significantly negative and red – significantly positive beta coefficients.

#### Key taxa representing major shift in the communities

To identify key taxa representing major shifts in the communities across the different treatments, differential abundance analysis was performed and the returned ASVs were an order of magnitude different between treatments. A larger log2 fold change is indicative of enrichment within each group. Common oil-degraders from the genera *Marinobacter*, *Oleispira*, and *Pseudomonas* were enriched in all treatments. *Vibrio*, *Oleiphilus,* and *Glaciecola* were enriched exclusively in CEWAF, while *Alcanivorax*, *Colwellia* and *Thalassotalea* (of the family *Colwelliaceae*) were enriched in both CEWAF and WAF. *Cycloclasticus*, *Pseudohongiella,* and *Acinetobacter* were enriched in the BEWAF and WAF treatments, whereas *Alteromonas*, *Moritella* and *Paraglaciecola* were enriched exclusively in the BEWAF (Supplementary File 5).

#### Discussion

##### Impact on microbial diversity

We observed lower species richness (alpha diversity) in treatments with added dispersant (CEWAF and SWD) compared to the BEWAF and WAF treatments. For the CEWAF and SWD treatments, this can be explained by the bloom of a few specialist taxa in the first seven days of incubation, which had responded directly to the dispersed crude oil or the dispersant itself. To further confirm the main factors that significantly altered the species richness and beta diversity in our treatments, we performed supervised machine learning modelling (regression analysis). The regression analysis showed that the presence of Finasol indeed negatively affected the species richness of the FSC bacterial community, and more so than exposure by the oil or it also with the rhamnolipid. If two biological systems share many characteristics (i.e. temperature, depth, nutrient availability, species etc.), data generated in one system can help inform what changes can be expected in another system for which limited data exist. While conventional analyses on microbiomes focus more on the variability of community structure in view of extrinsic environmental parameters, the regression analyses, on the other hand, can offer directionality (positive/negative contribution) to diversity analyses, and are widely useful to delineate diversity changes in multitude of environments. To our knowledge, supervised machine learning modelling has rarely been used in marine microbial metagenomic studies, especially in the context of hydrocarbon pollution and synthetic dispersant application. However, examples of using regression analyses in other microbial community studies exist [1]. It was interesting to note that the addition of crude oil alone (WAF) did not affect the alpha and beta diversity significantly, indicating that the microbial community in this treatment maintained a higher diversity over time.

**Table S1** Regression analysis top 10 models for Richness. The parameters of the model with the lowest Cross-validation error are given afterwards with significant positive influencers in red and negative in blue.

| | Model | Cross-validation Errors |
|---|---|---|
| M8 | Richness ~ Treatment_BEWAF + Treatment_CEWAF + Treatment_SWD + Treatment_SWBS + Treatment_FSC + Incubation_day_0 + Incubation_day_3 + Incubation_day_7 | 33.60742 |
| M7 | Richness ~ Treatment_BEWAF + Treatment_CEWAF + Treatment_SWD + Treatment_FSC + Incubation_day_0 + Incubation_day_3 + Incubation_day_7 | 33.70579 |
| M6 | Richness ~ Treatment_CEWAF + Treatment_SWD + Treatment_FSC + Incubation_day_0 + Incubation_day_3 + Incubation_day_7 | 34.72055 |
| M9 | Richness ~ Treatment_BEWAF + Treatment_CEWAF + Treatment_SWD + Treatment_SWBS + Treatment_FSC + Incubation_day_0 + Incubation_day_7 + Incubation_day_14 + Incubation_day_28 | 34.76996 |
| M10 | Richness ~ Treatment_WAF + Treatment_BEWAF + Treatment_CEWAF + Treatment_SWD + Treatment_SWBS + Treatment_FSC + Incubation_day_3 + Incubation_day_7 + Incubation_day_14 + Incubation_day_28 | 35.12272 |
| M5 | Richness ~ Treatment_CEWAF + Treatment_SWD + Incubation_day_0 + Incubation_day_3 + Incubation_day_7 | 37.53631 |
| M4 | Richness ~ Treatment_CEWAF + Treatment_SWD + Incubation_day_0 + Incubation_day_3 | 40.08586 |
| M3 | Richness ~ Treatment_SWD + Incubation_day_0 + Incubation_day_3 | 43.45814 |
| M2 | Richness ~ Treatment_SWD + Incubation_day_0 | 47.17362 |
| M1 | Richness ~ Incubation_day_0 | 52.23869 |

| Richness M8 | | | | | | | |
|---|---|---|---|---|---|---|---|
| **Predictors** | ***Estimates*** | ***std. Error*** | ***std. Beta*** | ***CI*** | ***standardized CI*** | ***Statistic*** | ***p*** |
| (Intercept) | 180.69595 *** | 7.59715 | | 165.58000 – 195.81190 | | 23.78470 | <0.001 |
| Treatment_BEWAF | 20.84874 | 10.68310 | 0.10368 | -0.40728 – 42.10476 | -0.00045 – 0.20781 | 1.95156 | 0.054 |
| Treatment_CEWAF | -39.09203 *** | 10.68310 | -0.19441 | -60.34805 – -17.83600 | -0.29854 – -0.09028 | -3.65924 | <0.001 |
| Treatment_SWD | -66.88116 *** | 11.23861 | -0.31374 | -89.24246 – -44.51985 | -0.41708 – -0.21041 | -5.95102 | <0.001 |
| Treatment_SWBS | -18.26529 | 10.68310 | -0.09084 | -39.52131 – 2.99074 | -0.19497 – 0.01329 | -1.70974 | 0.091 |
| Treatment_FSC | 79.71637 *** | 21.71171 | 0.19095 | 36.51688 – 122.91587 | 0.08902 – 0.29289 | 3.67158 | <0.001 |
| Incubation_day_0 | 92.50323 *** | 9.99698 | 0.51319 | 72.61238 – 112.39409 | 0.40449 – 0.62189 | 9.25312 | <0.001 |
| Incubation_day_3 | -62.37200 *** | 9.80121 | -0.33293 | -81.87332 – -42.87067 | -0.43546 – -0.23039 | -6.36371 | <0.001 |
| Incubation_day_7 | -38.28133 *** | 9.80121 | -0.20434 | -57.78265 – -18.78000 | -0.30687 – -0.10180 | -3.90578 | <0.001 |
| **Observations** | 90 | | | | | | |
| **$R^2$ / $R^2$ adjusted** | 0.819 / 0.801 | | | | | | |
| | | | | | | | *** p<0.05 ** p<0.01 *** p<0.001** |

**Table S2** Regression analysis top 10 models for Shannon index. The parameters of the model with the lowest Cross-validation error are given afterwards with significant positive influencers in red and negative in blue.

| | Model | Cross-validation Errors |
|---|---|---|
| M6 | Shannon ~ Treatment_BEWAF + Treatment_SWD + Treatment_SWBS + Incubation_day_0 + Incubation_day_3 + Incubation_day_28 | 0.52981 |
| M5 | Shannon ~ Treatment_BEWAF + Treatment_SWD + Incubation_day_0 + Incubation_day_3 + Incubation_day_28 | 0.53067 |
| M4 | Shannon ~ Treatment_BEWAF + Treatment_SWD + Incubation_day_0 + Incubation_day_3 | 0.53191 |
| M7 | Shannon ~ Treatment_BEWAF + Treatment_SWD + Treatment_SWBS + Incubation_day_0 + Incubation_day_3 + Incubation_day_7 + Incubation_day_28 | 0.53729 |
| M9 | Shannon ~ Treatment_BEWAF + Treatment_CEWAF + Treatment_SWD + Treatment_SWBS + Treatment_SW + Incubation_day_0 + Incubation_day_3 + Incubation_day_7 + Incubation_day_14 | 0.53944 |
| M8 | Shannon ~ Treatment_BEWAF + Treatment_CEWAF + Treatment_SWD + Treatment_SWBS + Incubation_day_3 + Incubation_day_7 + Incubation_day_14 + Incubation_day_28 | 0.53996 |
| M10 | Shannon ~ Treatment_WAF + Treatment_BEWAF + Treatment_CEWAF + Treatment_SWD + Treatment_SWBS + Treatment_SW + Incubation_day_0 + Incubation_day_3 + Incubation_day_7 + Incubation_day_14 | 0.54886 |
| M3 | Shannon ~ Treatment_SWD + Incubation_day_0 + Incubation_day_3 | 0.55250 |
| M2 | Shannon ~ Treatment_SWD + Incubation_day_0 | 0.57346 |
| M1 | Shannon ~ Incubation_day_0 | 0.62367 |

| Shannon M6 | | | | | | | |
|---|---|---|---|---|---|---|---|
| ***Predictors*** | *Estimates* | *std. Error* | *std. Beta* | *CI* | *standardized CI* | *Statistic* | *p* |
| (Intercept) | 2.75576 *** | 0.10417 | | 2.54856 – 2.96296 | | 26.45324 | <0.001 |
| Treatment_BEWAF | 0.48462 ** | 0.15655 | 0.21540 | 0.17325 – 0.79598 | 0.07902 – 0.35178 | 3.09562 | 0.003 |
| Treatment_SWD | -0.53318 ** | 0.16627 | -0.22355 | -0.86389 – -0.20248 | -0.36018 – -0.08691 | -3.20670 | 0.002 |
| Treatment_SWBS | 0.16425 | 0.15655 | 0.07301 | -0.14712 – 0.47562 | -0.06337 – 0.20939 | 1.04920 | 0.297 |
| Incubation_day_0 | 1.31226 *** | 0.14727 | 0.65067 | 1.01936 – 1.60517 | 0.50755 – 0.79379 | 8.91081 | <0.001 |
| Incubation_day_3 | -0.41187 ** | 0.15233 | -0.19649 | -0.71484 – -0.10890 | -0.33892 – -0.05406 | -2.70389 | 0.008 |
| Incubation_day_28 | 0.25684 | 0.15941 | 0.11712 | -0.06022 – 0.57390 | -0.02535 – 0.25958 | 1.61119 | 0.111 |
| **Observations** | 90 | | | | | | |
| **$R^2$ / $R^2$ adjusted** | 0.635 / 0.608 | | | | | | |
| | | | | | | | *** p<0.05 ** p<0.01 *** p<0.001** |

**Table S3** Regression analysis top 10 models for NTI. The parameters of the model with the lowest Cross-validation error are given afterwards with significant positive influencers in red and negative in blue.

| | Model | Cross-validation Errors |
|---|---|---|
| **M6** | NTI ~ Treatment_CEWAF + Treatment_SWD + Treatment_SW + Treatment_FSC + Incubation_day_7 + Incubation_day_14 | 0.98634 |
| **M7** | NTI ~ Treatment_CEWAF + Treatment_SWD + Treatment_SWBS + Treatment_SW + Treatment_FSC + Incubation_day_7 + Incubation_day_14 | 0.98841 |
| **M8** | NTI ~ Treatment_CEWAF + Treatment_SWD + Treatment_SWBS + Treatment_SW + Treatment_FSC + Incubation_day_0 + Incubation_day_7 + Incubation_day_14 | 0.98919 |
| **M5** | NTI ~ Treatment_SWD + Treatment_SW + Treatment_FSC + Incubation_day_7 + Incubation_day_14 | 0.99991 |
| **M9** | NTI ~ Treatment_WAF + Treatment_BEWAF + Treatment_CEWAF + Treatment_SWBS + Treatment_FSC + Incubation_day_3 + Incubation_day_7 + Incubation_day_14 + Incubation_day_28 | 1.00014 |
| **M4** | NTI ~ Treatment_SWD + Treatment_SW + Treatment_FSC + Incubation_day_7 | 1.00617 |
| **M10** | NTI ~ Treatment_WAF + Treatment_CEWAF + Treatment_SWD + Treatment_SWBS + Treatment_SW + Treatment_FSC + Incubation_day_0 + Incubation_day_3 + Incubation_day_7 + Incubation_day_28 | 1.01131 |
| **M3** | NTI ~ Treatment_CEWAF + Treatment_FSC + Incubation_day_7 | 1.01589 |
| **M2** | NTI ~ Treatment_FSC + Incubation_day_7 | 1.05334 |
| **M1** | NTI ~ Treatment_FSC | 1.08783 |

| NTI M6 | | | | | | | |
|---|---|---|---|---|---|---|---|
| *Predictors* | *Estimates* | *std. Error* | *std. Beta* | *CI* | *standardized CI* | *Statistic* | *p* |
| (Intercept) | 6.58923 *** | 0.17336 | | 6.24443 – 6.93402 | | 38.00993 | <0.001 |
| Treatment_CEWAF | 0.48364 | 0.29596 | 0.15687 | -0.10501 – 1.07230 | -0.03128 – 0.34501 | 1.63414 | 0.106 |
| Treatment_SWD | -0.64640 * | 0.31268 | -0.19776 | -1.26830 – -0.02450 | -0.38526 – -0.01027 | -2.06730 | 0.042 |
| Treatment_SW | -0.69154 * | 0.29596 | -0.22430 | -1.28020 – -0.10289 | -0.41244 – -0.03615 | -2.33659 | 0.022 |
| Treatment_FSC | -1.90112 ** | 0.59721 | -0.29700 | -3.08894 – -0.71329 | -0.47986 – -0.11414 | -3.18334 | 0.002 |
| Incubation_day_7 | 0.88635 ** | 0.27153 | 0.30856 | 0.34630 – 1.42640 | 0.12329 – 0.49382 | 3.26433 | 0.002 |
| Incubation_day_14 | 0.46463 | 0.27153 | 0.16175 | -0.07542 – 1.00469 | -0.02352 – 0.34701 | 1.71119 | 0.091 |
| **Observations** | 90 | | | | | | |
| **$R^2$ / $R^2$ adjusted** | 0.316 / 0.266 | | | | | | |
| | | | | | | | *** p<0.05 ** p<0.01 *** p<0.001** |

**Table S4** Regression analysis top 10 models for NRI. The parameters of the model with the lowest Cross-validation error are given afterwards with significant positive influencers in red and negative in blue.

| | Model | Cross-validation Errors |
|---|---|---|
| **M8** | NRI ~ Treatment_WAF + Treatment_BEWAF + Treatment_CEWAF + Treatment_SWD + Treatment_SWBS + Incubation_day_0 + Incubation_day_3 + Incubation_day_14 | 1.11824 |
| **M7** | NRI ~ Treatment_BEWAF + Treatment_CEWAF + Treatment_SWD + Treatment_SW + Incubation_day_0 + Incubation_day_3 + Incubation_day_14 | 1.11853 |
| **M9** | NRI ~ Treatment_WAF + Treatment_BEWAF + Treatment_CEWAF + Treatment_SWD + Treatment_SWBS + Treatment_SW + Incubation_day_0 + Incubation_day_3 + Incubation_day_14 | 1.12987 |
| **M6** | NRI ~ Treatment_BEWAF + Treatment_CEWAF + Treatment_SWD + Treatment_SW + Incubation_day_0 + Incubation_day_3 | 1.13572 |
| **M5** | NRI ~ Treatment_BEWAF + Treatment_CEWAF + Treatment_SWD + Incubation_day_0 + Incubation_day_3 | 1.14052 |
| **M10** | NRI ~ Treatment_WAF + Treatment_BEWAF + Treatment_CEWAF + Treatment_SWD + Treatment_SWBS + Treatment_FSC + Incubation_day_3 + Incubation_day_7 + Incubation_day_14 + Incubation_day_28 | 1.14529 |
| **M4** | NRI ~ Treatment_CEWAF + Treatment_SWD + Incubation_day_0 + Incubation_day_3 | 1.19900 |
| **M3** | NRI ~ Treatment_CEWAF + Treatment_SWD + Incubation_day_0 | 1.27369 |
| **M2** | NRI ~ Treatment_SWD + Incubation_day_0 | 1.54889 |
| **M1** | NRI ~ Incubation_day_0 | 1.70878 |

| **NRI M8** | | | | | | | |
|---|---|---|---|---|---|---|---|
| *Predictors* | *Estimates* | *std. Error* | *std. Beta* | *CI* | *standardized CI* | *Statistic* | *p* |
| (Intercept) | 1.81273 *** | 0.29594 | | 1.22390 – 2.40157 | | 6.12525 | <0.001 |
| Treatment_WAF | 0.88162 * | 0.38237 | 0.14220 | 0.12083 – 1.64241 | 0.02132 – 0.26308 | 2.30570 | 0.024 |
| Treatment_BEWAF | 1.61687 *** | 0.37343 | 0.26817 | 0.87387 – 2.35988 | 0.14678 – 0.38956 | 4.32983 | <0.001 |
| Treatment_CEWAF | 3.26464 *** | 0.37343 | 0.54146 | 2.52164 – 4.00764 | 0.42007 – 0.66285 | 8.74239 | <0.001 |
| Treatment_SWD | 3.45907 *** | 0.38842 | 0.54117 | 2.68624 – 4.23190 | 0.42206 – 0.66027 | 8.90555 | <0.001 |
| Treatment_SWBS | 0.72814 | 0.37343 | 0.12077 | -0.01486 – 1.47114 | -0.00062 – 0.24216 | 1.94988 | 0.055 |
| Incubation_day_0 | -3.28898 *** | 0.30243 | -0.60853 | -3.89072 – -2.68725 | -0.71820 – -0.49886 | -10.87535 | <0.001 |
| Incubation_day_3 | 0.85700 ** | 0.31027 | 0.15256 | 0.23966 – 1.47435 | 0.04430 – 0.26082 | 2.76209 | 0.007 |
| Incubation_day_14 | -0.56575 | 0.31027 | -0.10071 | -1.18310 – 0.05160 | -0.20897 – 0.00754 | -1.82339 | 0.072 |
| **Observations** | 90 | | | | | | |
| **$R^2$ / $R^2$ adjusted** | 0.799 / 0.779 | | | | | | |
| | | | | | | | *** p<0.05 ** p<0.01 *** p<0.001** |

**Table S5** Regression analysis top 10 models for LCBD (Bray-Curtis). The parameters of the model with the lowest Cross-validation error are given afterwards with significant positive influencers in red and negative in blue.

| | Model | Cross-validation Errors |
|---|---|---|
| M8 | LCBD ~ Treatment_WAF + Treatment_CEWAF + Treatment_SWD + Treatment_SW + Treatment_FSC + Incubation_day_0 + Incubation_day_14 + Incubation_day_28 | 0.00102 |
| M9 | LCBD ~ Treatment_WAF + Treatment_CEWAF + Treatment_SWD + Treatment_SW + Treatment_FSC + Incubation_day_0 + Incubation_day_3 + Incubation_day_7 + Incubation_day_28 | 0.00103 |
| M10 | LCBD ~ Treatment_WAF + Treatment_BEWAF + Treatment_CEWAF + Treatment_SWD + Treatment_SWBS + Treatment_SW + Incubation_day_0 + Incubation_day_3 + Incubation_day_7 + Incubation_day_28 | 0.00106 |
| M7 | LCBD ~ Treatment_BEWAF + Treatment_SWD + Treatment_SWBS + Treatment_FSC + Incubation_day_0 + Incubation_day_14 + Incubation_day_28 | 0.00108 |
| M6 | LCBD ~ Treatment_BEWAF + Treatment_SWD + Treatment_SWBS + Treatment_FSC + Incubation_day_14 + Incubation_day_28 | 0.00115 |
| M5 | LCBD ~ Treatment_BEWAF + Treatment_CEWAF + Treatment_SWBS + Incubation_day_14 + Incubation_day_28 | 0.00119 |
| M4 | LCBD ~ Treatment_BEWAF + Treatment_SWBS + Incubation_day_14 + Incubation_day_28 | 0.00127 |
| M3 | LCBD ~ Treatment_BEWAF + Treatment_SWBS + Incubation_day_28 | 0.00138 |
| M2 | LCBD ~ Treatment_BEWAF + Incubation_day_28 | 0.00157 |
| M1 | LCBD ~ Incubation_day_28 | 0.00169 |

| LCBD Bray-Curtis M8 | | | | | | | |
|---|---|---|---|---|---|---|---|
| *Predictors* | *Estimates* | *std. Error* | *std. Beta* | *CI* | *standardized CI* | *Statistic* | *p* |
| (Intercept) | 0.00915 *** | 0.00022 | | 0.00871 – 0.00959 | | 41.47606 | **<0.001** |
| Treatment_WAF | 0.00219 *** | 0.00032 | 0.40460 | 0.00156 – 0.00283 | 0.28952 – 0.51968 | 6.89090 | **<0.001** |
| Treatment_SWD | 0.00301 *** | 0.00033 | 0.53837 | 0.00236 – 0.00366 | 0.42338 – 0.65335 | 9.17646 | **0.001** |
| Treatment_CEWAF | 0.00112 *** | 0.00031 | 0.21336 | 0.00051 – 0.00174 | 0.09789 – 0.32884 | 3.62132 | **<0.001** |
| Treatment_SW | 0.00178 *** | 0.00031 | 0.33850 | 0.00117 – 0.00240 | 0.22302 – 0.45398 | 5.74523 | **<0.001** |
| Treatment_FSC | 0.00489 *** | 0.00063 | 0.44685 | 0.00363 – 0.00615 | 0.33372 – 0.55998 | 7.74154 | **<0.001** |
| Incubation_day_0 | -0.00115 *** | 0.00029 | -0.24326 | -0.00172 – -0.00057 | -0.36328 – -0.12323 | -3.97233 | **<0.001** |
| Incubation_day_14 | 0.00127 *** | 0.00028 | -0.25957 | 0.00071 – 0.00184 | 0.14642 – 0.37272 | 4.49640 | **<0.001** |
| Incubation_day_28 | 0.00304 *** | 0.00030 | 0.59274 | 0.00245 – 0.00364 | 0.47955 – 0.70592 | 10.26445 | **<0.001** |
| **Observations** | 90 | | | | | | |
| **$R^2$ / $R^2$ adjusted** | 0.775 / 0.753 | | | | | | |
| | | | | | | | *** p<0.05 ** p<0.01 *** p<0.001** |

**Table S6** Regression analysis top 10 models for LCBD (unweighted UniFrac). The parameters of the model with the lowest Cross-validation error are given afterwards with significant positive influencers in red and negative in blue.

| | Model | Cross-validation Errors |
|---|---|---|
| M7 | LCBD ~ Treatment_WAF + Treatment_SWD + Treatment_SW + Treatment_FSC + Incubation_day_3 + Incubation_day_7 + Incubation_day_28 | 0.00083 |
| M8 | LCBD ~ Treatment_WAF + Treatment_BEWAF + Treatment_SWD + Treatment_SW + Treatment_FSC + Incubation_day_3 + Incubation_day_7 + Incubation_day_28 | 0.00083 |
| M6 | LCBD ~ Treatment_SWD + Treatment_SW + Treatment_FSC + Incubation_day_3 + Incubation_day_7 + Incubation_day_28 | 0.00084 |
| M9 | LCBD ~ Treatment_WAF + Treatment_BEWAF + Treatment_SWD + Treatment_SW + Treatment_FSC + Incubation_day_0 + Incubation_day_3 + Incubation_day_7 + Incubation_day_28 | 0.00084 |
| M4 | LCBD ~ Treatment_SWD + Treatment_FSC + Incubation_day_3 + Incubation_day_28 | 0.00085 |
| M5 | LCBD ~ Treatment_SWD + Treatment_FSC + Incubation_day_3 + Incubation_day_7 + Incubation_day_28 | 0.00085 |
| M10 | LCBD ~ Treatment_WAF + Treatment_BEWAF + Treatment_SWD + Treatment_SWBS + Treatment_SW + Treatment_FSC + Incubation_day_0 + Incubation_day_3 + Incubation_day_7 + Incubation_day_14 | 0.00085 |
| M3 | LCBD ~ Treatment_SWD + Treatment_FSC + Incubation_day_28 | 0.00093 |
| M2 | LCBD ~ Treatment_FSC + Incubation_day_28 | 0.00105 |
| M1 | LCBD ~ Incubation_day_28 | 0.00119 |

**LCBD Unweighted UniFrac M7**

| *Predictors* | *Estimates* | *std. Error* | *std. Beta* | *CI* | *standardized CI* | *Statistic* | *p* |
|---|---|---|---|---|---|---|---|
| (Intercept) | 0.01071 *** | 0.00015 | | 0.01040 – 0.01101 | | 70.10001 | <0.001 |
| Treatment_WAF | 0.00044 | 0.00024 | 0.11977 | -0.00003 – 0.00090 | -0.00670 – 0.24623 | 1.85613 | 0.067 |
| Treatment_SWD | 0.00174 *** | 0.00024 | 0.46183 | 0.00125 – 0.00222 | 0.33493 – 0.58873 | 7.13305 | <0.001 |
| Treatment_SW | 0.00066 ** | 0.00023 | 0.18508 | 0.00020 – 0.00111 | 0.05845 – 0.31171 | 2.86468 | 0.005 |
| Treatment_FSC | 0.00307 *** | 0.00047 | 0.41684 | 0.00214 – 0.00400 | 0.29192 – 0.54176 | 6.54018 | <0.001 |
| Incubation_day_3 | -0.00125 *** | 0.00022 | -0.37971 | -0.00170 – -0.00081 | -0.51197 – -0.24745 | -5.62684 | <0.001 |
| Incubation_day_7 | -0.00060 ** | 0.00022 | -0.18051 | -0.00104 – -0.00015 | -0.31277 – -0.04825 | -2.67496 | 0.009 |
| Incubation_day_28 | 0.00136 *** | 0.00023 | 0.39479 | 0.00090 – 0.00183 | 0.26248 – 0.52710 | 5.84801 | <0.001 |
| **Observations** | 90 | | | | | | |
| **$R^2$ / $R^2$ adjusted** | 0.692 / 0.666 | | | | | | |

*** p<0.05 ** p<0.01 *** p<0.001**

**Table S7** Regression analysis top 10 models for LCBD (weighted UniFrac). The parameters of the model with the lowest Cross-validation error are given afterwards with significant positive influencers in red and negative in blue.

| | Model | Cross-validation Errors |
|---|---|---|
| M8 | LCBD ~ Treatment_WAF + Treatment_CEWAF + Treatment_SWD + Treatment_SWBS + Treatment_FSC + Incubation_day_7 + Incubation_day_14 + Incubation_day_28 | 0.00449 |
| M7 | LCBD ~ Treatment_WAF + Treatment_CEWAF + Treatment_SWD + Treatment_FSC + Incubation_day_7 + Incubation_day_14 + Incubation_day_28 | 0.00450 |
| M6 | LCBD ~ Treatment_WAF + Treatment_CEWAF + Treatment_SWD + Treatment_FSC + Incubation_day_14 + Incubation_day_28 | 0.00454 |
| M9 | LCBD ~ Treatment_WAF + Treatment_BEWAF + Treatment_CEWAF + Treatment_SWD + Treatment_SWBS + Treatment_SW + Incubation_day_7 + Incubation_day_14 + Incubation_day_28 | 0.00455 |
| M10 | LCBD ~ Treatment_WAF + Treatment_BEWAF + Treatment_CEWAF + Treatment_SWBS + Treatment_SW + Treatment_FSC + Incubation_day_0 + Incubation_day_3 + Incubation_day_14 + Incubation_day_28 | 0.00455 |
| M5 | LCBD ~ Treatment_CEWAF + Treatment_SWD + Treatment_FSC + Incubation_day_14 + Incubation_day_28 | 0.00460 |
| M4 | LCBD ~ Treatment_CEWAF + Treatment_SWD + Treatment_FSC + Incubation_day_28 | 0.00461 |
| M3 | LCBD ~ Treatment_CEWAF + Treatment_SWD + Treatment_FSC | 0.00481 |
| M2 | LCBD ~ Treatment_SWD + Treatment_FSC | 0.00508 |
| M1 | LCBD ~ Treatment_SWD | 0.00555 |

| **LCBD Weighted UniFrac M8** | | | | | | | |
|---|---|---|---|---|---|---|---|
| *Predictors* | *Estimates* | *std. Error* | *std. Beta* | *CI* | *standardized CI* | *Statistic* | *p* |
| (Intercept) | 0.00627 *** | 0.00097 | | 0.00434 – 0.00820 | | 6.47096 | <0.001 |
| Treatment_WAF | 0.00176 | 0.00139 | 0.10139 | -0.00102 – 0.00453 | -0.05630 – 0.25908 | 1.26016 | 0.211 |
| Treatment_CEWAF | 0.00529 *** | 0.00136 | 0.31377 | 0.00258 – 0.00800 | 0.15542 – 0.47213 | 3.88362 | <0.001 |
| Treatment_SWD | 0.01063 *** | 0.00144 | 0.59516 | 0.00777 – 0.01349 | 0.43748 – 0.75284 | 7.39793 | <0.001 |
| Treatment_SWBS | -0.00157 | 0.00136 | -0.09306 | -0.00428 – 0.00114 | -0.25142 – 0.06529 | -1.15186 | 0.253 |
| Treatment_FSC | 0.01631 *** | 0.00267 | 0.46608 | 0.01100 – 0.02162 | 0.31662 – 0.61554 | 6.11212 | <0.001 |
| Incubation_day_7 | 0.00149 | 0.00125 | 0.09473 | -0.00100 – 0.00397 | -0.06120 – 0.25067 | 1.19072 | 0.237 |
| Incubation_day_14 | 0.00378 ** | 0.00125 | 0.24070 | 0.00129 – 0.00626 | 0.08476 – 0.39664 | 3.02537 | 0.003 |
| Incubation_day_28 | 0.00459 *** | 0.00131 | 0.27923 | 0.00199 – 0.00719 | 0.12323 – 0.43522 | 3.50822 | 0.001 |
| **Observations** | 90 | | | | | | |
| **$R^2$ / $R^2$ adjusted** | 0.577 / 0.535 | | | | | | |
| | | | | | | | *** p<0.05 ** p<0.01 *** p<0.001** |

**Table S8.** Statistical summary of all predicted pathways which were significantly different between treatments BEWAF, CEWAF, and WAF (ANOVA). Size means the total KEGG orthologs (KO) that are present in the particular pathway. Hits is the number of KEGG orthologs that were predicted in our data set. Pval is p-value based on ANOVA and FDR is the false discovery rate. The table was generated in Microbiomeanalyst online tool [2] using the KO abundance table generated in PICRUsT2.

| Pathway | Size | Hits | Statistic.Q | Expected.Q | Pval | FDR |
|---|---|---|---|---|---|---|
| Carotenoid biosynthesis | 27 | 15 | 9.97168466 | 1.13596366 | 1.94E-12 | 2.68E-10 |
| Xylene degradation | 20 | 18 | 5.28809519 | 1.13591273 | 4.16E-10 | 2.14E-08 |
| Betalain biosynthesis | 3 | 3 | 9.12897169 | 1.13595138 | 6.12E-10 | 2.14E-08 |
| Indole alkaloid biosynthesis | 3 | 1 | 9.13653965 | 1.13595062 | 6.19E-10 | 2.14E-08 |
| Degradation of aromatic compounds | 134 | 81 | 6.18544513 | 1.1361019 | 3.05E-09 | 8.41E-08 |
| Styrene degradation | 12 | 9 | 6.34546 | 1.13611384 | 1.17E-08 | 2.69E-07 |
| Glycosylphosphatidylinositol(GPI)-anchor biosynthesis | 17 | 1 | 5.40109822 | 1.13488291 | 2.81E-08 | 5.54E-07 |
| Toluene degradation | 35 | 24 | 4.40880673 | 1.13598138 | 1.32E-07 | 2.28E-06 |
| Penicillin and cephalosporin biosynthesis | 8 | 2 | 6.57054924 | 1.1356708 | 2.71E-07 | 4.16E-06 |
| Dioxin degradation | 25 | 18 | 4.15338553 | 1.13604411 | 5.49E-07 | 7.06E-06 |
| Phosphonate and phosphinate metabolism | 9 | 7 | 4.74904 | 1.13602909 | 5.63E-07 | 7.06E-06 |
| Chlorocyclohexane and chlorobenzene degradation | 37 | 22 | 5.18354963 | 1.13608511 | 1.42E-06 | 1.64E-05 |
| Fatty acid elongation | 9 | 4 | 4.11550624 | 1.13578324 | 3.71E-06 | 3.76E-05 |
| Atrazine degradation | 10 | 7 | 6.22756395 | 1.13610806 | 3.84E-06 | 3.76E-05 |
| Ethylbenzene degradation | 13 | 10 | 5.06309761 | 1.13593171 | 4.08E-06 | 3.76E-05 |
| Polycyclic aromatic hydrocarbon degradation | 30 | 11 | 5.3210803 | 1.13605253 | 4.48E-06 | 3.87E-05 |
| Brassinosteroid biosynthesis | 8 | 1 | 5.83073314 | 1.13599469 | 1.33E-05 | 0.00010813 |
| Sphingolipid metabolism | 29 | 9 | 5.38022467 | 1.1361375 | 1.72E-05 | 0.00013151 |
| Nicotinate and nicotinamide metabolism | 36 | 24 | 4.26285119 | 1.13616091 | 2.18E-05 | 0.00015039 |
| Galactose metabolism | 56 | 36 | 3.97106009 | 1.13616143 | 2.18E-05 | 0.00015039 |
| Sulfur metabolism | 38 | 33 | 4.35511156 | 1.1361461 | 2.89E-05 | 0.00018992 |
| Steroid biosynthesis | 26 | 12 | 2.6769289 | 1.13568296 | 3.85E-05 | 0.00024157 |
| Phenylpropanoid biosynthesis | 18 | 7 | 4.20373168 | 1.13615719 | 4.89E-05 | 0.00029369 |
| Other glycan degradation | 7 | 6 | 4.26344332 | 1.1361368 | 6.28E-05 | 0.00036083 |
| Isoquinoline alkaloid biosynthesis | 34 | 8 | 3.96347065 | 1.13616444 | 9.80E-05 | 0.00054072 |
| Glycine, serine and threonine metabolism | 78 | 68 | 3.9838976 | 1.13614447 | 0.00013135 | 0.00069716 |
| Porphyrin and chlorophyll metabolism | 69 | 61 | 3.78837438 | 1.13616769 | 0.00015668 | 0.00080079 |
| Chloroalkane and chloroalkene degradation | 23 | 18 | 4.43373845 | 1.13611423 | 0.00016906 | 0.00083323 |
| Ubiquinone and other terpenoid-quinone biosynthesis | 40 | 32 | 3.67440507 | 1.13616153 | 0.00018231 | 0.00086753 |

| | | | | | | |
|---|---|---|---|---|---|---|
| Pentose and glucuronate interconversions | 41 | 29 | 4.34074766 | 1.13612532 | 0.00019185 | 0.00088252 |
| Thiamine metabolism | 23 | 19 | 3.64862103 | 1.13617186 | 0.00022895 | 0.00100231 |
| Ascorbate and aldarate metabolism | 31 | 21 | 4.59681155 | 1.13610289 | 0.00023242 | 0.00100231 |
| Fructose and mannose metabolism | 44 | 39 | 3.51012366 | 1.136164 | 0.00025099 | 0.0010496 |
| alpha-Linolenic acid metabolism | 18 | 3 | 4.61799267 | 1.13596199 | 0.00032437 | 0.00131654 |
| Arginine and proline metabolism | 115 | 87 | 3.77211405 | 1.13614983 | 0.00037629 | 0.00148366 |
| Fluorobenzoate degradation | 11 | 10 | 3.48659567 | 1.13601335 | 0.00043032 | 0.00164958 |
| Glycerolipid metabolism | 57 | 21 | 4.01485804 | 1.13613582 | 0.00044418 | 0.00165667 |
| Histidine metabolism | 35 | 29 | 3.87528775 | 1.13614082 | 0.00045897 | 0.00166678 |
| Tyrosine metabolism | 54 | 29 | 3.545479 | 1.13616393 | 0.00051483 | 0.00182172 |
| Taurine and hypotaurine metabolism | 17 | 14 | 3.34317642 | 1.1361523 | 0.00061986 | 0.00213851 |
| Ether lipid metabolism | 26 | 6 | 4.00333279 | 1.13607104 | 0.00066002 | 0.00222153 |
| Limonene and pinene degradation | 7 | 5 | 4.00686728 | 1.13611285 | 0.00068264 | 0.00224296 |
| Arachidonic acid metabolism | 42 | 2 | 4.41942989 | 1.1360507 | 0.00073724 | 0.00236603 |
| Flavonoid biosynthesis | 12 | 2 | 4.21685835 | 1.13584686 | 0.00081901 | 0.00256873 |
| Cysteine and methionine metabolism | 71 | 55 | 3.27883171 | 1.13615863 | 0.0008919 | 0.00273517 |
| Folate biosynthesis | 29 | 25 | 3.2721364 | 1.13616078 | 0.0009636 | 0.00289081 |
| Selenocompound metabolism | 15 | 13 | 3.43120091 | 1.13616047 | 0.00111601 | 0.0032768 |
| Alanine, aspartate and glutamate metabolism | 62 | 47 | 3.34549003 | 1.1361575 | 0.0012383 | 0.00356011 |
| Benzoate degradation | 59 | 46 | 3.37205583 | 1.1361471 | 0.00136426 | 0.0038422 |
| Starch and sucrose metabolism | 65 | 50 | 2.94813513 | 1.13616902 | 0.00153493 | 0.00423641 |
| Methane metabolism | 105 | 72 | 3.16793163 | 1.13616318 | 0.00158714 | 0.00429461 |
| beta-Alanine metabolism | 36 | 27 | 3.46033817 | 1.13612818 | 0.00176144 | 0.00467459 |
| Pentose phosphate pathway | 63 | 45 | 2.98081081 | 1.1361657 | 0.00187516 | 0.0048825 |
| Drug metabolism - other enzymes | 20 | 14 | 3.2831644 | 1.13614947 | 0.00200104 | 0.00511376 |
| Steroid degradation | 9 | 9 | 3.35937159 | 1.1360578 | 0.00221512 | 0.00555793 |
| Naphthalene degradation | 17 | 12 | 2.94100972 | 1.13613197 | 0.00225706 | 0.00556203 |
| Butirosin and neomycin biosynthesis | 3 | 2 | 3.88950241 | 1.13603624 | 0.00234952 | 0.00565521 |
| Cyanoamino acid metabolism | 10 | 7 | 3.23158042 | 1.13613768 | 0.00237683 | 0.00565521 |
| Tetracycline biosynthesis | 6 | 6 | 3.85238356 | 1.13607273 | 0.00247895 | 0.00579111 |
| Lipopolysaccharide biosynthesis | 17 | 17 | 3.48968071 | 1.13612783 | 0.002519 | 0.00579111 |
| Glycolysis / Gluconeogenesis | 80 | 62 | 3.1556835 | 1.13615494 | 0.00255984 | 0.00579111 |
| Drug metabolism - cytochrome P450 | 19 | 5 | 2.94129609 | 1.13612488 | 0.00268369 | 0.00597337 |
| Nitrotoluene degradation | 7 | 7 | 3.4639434 | 1.13594147 | 0.00275845 | 0.00604232 |
| Purine metabolism | 181 | 118 | 3.1452387 | 1.13615205 | 0.00339601 | 0.00719095 |
| One carbon pool by folate | 24 | 18 | 3.28647671 | 1.13611624 | 0.00343905 | 0.00719095 |
| Amino sugar and nucleotide sugar metabolism | 64 | 52 | 3.01190304 | 1.13615415 | 0.00343915 | 0.00719095 |
| Glycerophospholipid metabolism | 65 | 28 | 2.99047739 | 1.13615929 | 0.00467197 | 0.00962286 |
| Aminobenzoate degradation | 20 | 13 | 2.62855792 | 1.13613028 | 0.00491588 | 0.00997635 |

| | | | | | | |
|---|---|---|---|---|---|---|
| Phenylalanine, tyrosine and tryptophan biosynthesis | 64 | 50 | 2.93225118 | 1.13614178 | 0.00588982 | 0.01177965 |
| Pyrimidine metabolism | 149 | 92 | 3.00625875 | 1.13614791 | 0.0067611 | 0.01332903 |
| Pyruvate metabolism | 74 | 64 | 2.88508525 | 1.13615762 | 0.00726018 | 0.01411133 |
| Steroid hormone biosynthesis | 33 | 5 | 2.98297478 | 1.13593603 | 0.00761162 | 0.01458894 |
| Glutathione metabolism | 28 | 16 | 2.8467084 | 1.13614829 | 0.0098772 | 0.01841114 |
| Sesquiterpenoid and triterpenoid biosynthesis | 11 | 3 | 2.78815187 | 1.13518136 | 0.01023049 | 0.01841114 |
| Lysine degradation | 30 | 22 | 2.8702454 | 1.13614015 | 0.01023455 | 0.01841114 |
| Glycosaminoglycan biosynthesis - chondroitin sulfate / dermatan sulfate | 10 | 2 | 2.70286248 | 1.13561504 | 0.01035069 | 0.01841114 |
| Glycosaminoglycan biosynthesis - heparan sulfate / heparin | 9 | 2 | 2.70286248 | 1.13561504 | 0.01035069 | 0.01841114 |
| Retinol metabolism | 41 | 7 | 2.60807976 | 1.1361166 | 0.0105058 | 0.01841114 |
| Metabolism of xenobiotics by cytochrome P450 | 21 | 4 | 2.60795655 | 1.13611644 | 0.01053971 | 0.01841114 |
| Nitrogen metabolism | 17 | 16 | 2.92958808 | 1.13613307 | 0.01067356 | 0.0184119 |
| Inositol phosphate metabolism | 39 | 13 | 2.77623819 | 1.13614249 | 0.01119923 | 0.01908017 |
| Tryptophan metabolism | 42 | 24 | 2.80704319 | 1.13613961 | 0.01152336 | 0.01939296 |
| Carbapenem biosynthesis | 2 | 2 | 3.08588892 | 1.13610558 | 0.01297213 | 0.02156811 |
| Peptidoglycan biosynthesis | 13 | 10 | 3.01916776 | 1.13611262 | 0.01422744 | 0.02337365 |
| Carbon metabolism | 249 | 177 | 2.55921016 | 1.13616162 | 0.01516148 | 0.0246151 |
| Phenylalanine metabolism | 41 | 25 | 2.45206162 | 1.13616579 | 0.01601087 | 0.02560782 |
| Caprolactam degradation | 13 | 11 | 2.34638479 | 1.13614838 | 0.01614406 | 0.02560782 |
| Fatty acid degradation | 39 | 24 | 2.60667707 | 1.13614359 | 0.01680699 | 0.02635641 |
| Glycosaminoglycan degradation | 14 | 10 | 2.65700099 | 1.13611735 | 0.01746633 | 0.02708263 |
| Carbon fixation in photosynthetic organisms | 35 | 24 | 2.55865784 | 1.13615138 | 0.01878063 | 0.02879696 |
| Biosynthesis of amino acids | 222 | 195 | 2.54177795 | 1.13615116 | 0.01920682 | 0.02885052 |
| Valine, leucine and isoleucine degradation | 58 | 41 | 2.57720101 | 1.13614163 | 0.01950582 | 0.02885052 |
| Glyoxylate and dicarboxylate metabolism | 51 | 41 | 2.53849201 | 1.13615598 | 0.01958665 | 0.02885052 |
| D-Glutamine and D-glutamate metabolism | 6 | 6 | 2.57760322 | 1.13614301 | 0.01965181 | 0.02885052 |
| Novobiocin biosynthesis | 12 | 9 | 2.40828911 | 1.13616563 | 0.02026353 | 0.02943544 |
| Stilbenoid, diarylheptanoid and gingerol biosynthesis | 5 | 2 | 2.71013396 | 1.13556871 | 0.02110049 | 0.03033196 |
| Oxidative phosphorylation | 16 | 10 | 2.7380459 | 1.1361017 | 0.02272129 | 0.03232514 |
| Citrate cycle (TCA cycle) | 53 | 38 | 2.52373591 | 1.13615915 | 0.02320847 | 0.03268132 |
| Vitamin B6 metabolism | 12 | 9 | 2.52235243 | 1.13614189 | 0.02357859 | 0.03274728 |
| Pantothenate and CoA biosynthesis | 29 | 24 | 2.55685307 | 1.13614003 | 0.02372992 | 0.03274728 |
| Lysine biosynthesis | 41 | 35 | 2.48442961 | 1.13615221 | 0.02406931 | 0.03288677 |
| Geraniol degradation | 6 | 6 | 2.17660482 | 1.1361301 | 0.02679093 | 0.03624656 |
| Biosynthesis of siderophore group nonribosomal peptides | 3 | 2 | 2.33099871 | 1.13600902 | 0.02914228 | 0.03904499 |
| Glycosphingolipid biosynthesis - globo series | 10 | 1 | 2.65051469 | 1.13609443 | 0.0323611 | 0.04253174 |

| | | | | | | |
|---|---|---|---|---|---|---|
| Glycosphingolipid biosynthesis - ganglio series | 13 | 1 | 2.65051469 | 1.13609443 | 0.0323611 | 0.04253174 |
| Caffeine metabolism | 9 | 3 | 2.44282322 | 1.13549754 | 0.03342054 | 0.04350976 |
| Various types of N-glycan biosynthesis | 28 | 4 | 2.50795012 | 1.13593671 | 0.0353728 | 0.04562099 |
| Carbon fixation pathways in prokaryotes | 60 | 49 | 2.29028591 | 1.1361594 | 0.03849426 | 0.04918711 |
| Fatty acid biosynthesis | 28 | 23 | 2.31768643 | 1.13614242 | 0.04036355 | 0.05067933 |
| Linoleic acid metabolism | 20 | 3 | 2.19093346 | 1.13616274 | 0.04039657 | 0.05067933 |
| 2-Oxocarboxylic acid metabolism | 57 | 44 | 2.27845284 | 1.1361506 | 0.04155126 | 0.05165833 |
| Primary bile acid biosynthesis | 14 | 3 | 2.24930887 | 1.13610288 | 0.04376648 | 0.05392655 |
| C5-Branched dibasic acid metabolism | 11 | 11 | 2.33328086 | 1.1361248 | 0.04762938 | 0.05816686 |
| Flavone and flavonol biosynthesis | 3 | 1 | 2.43506139 | 1.13608911 | 0.04868892 | 0.05893921 |
| Biotin metabolism | 19 | 17 | 2.18432575 | 1.13614387 | 0.05213289 | 0.06255947 |
| Riboflavin metabolism | 22 | 14 | 2.22311294 | 1.1361467 | 0.05386956 | 0.0640862 |
| D-Alanine metabolism | 3 | 3 | 2.27691485 | 1.13614682 | 0.05485247 | 0.06469778 |
| Tropane, piperidine and pyridine alkaloid biosynthesis | 19 | 9 | 2.0954149 | 1.13616168 | 0.05804609 | 0.06788441 |
| Lipoic acid metabolism | 4 | 3 | 2.27358553 | 1.13613219 | 0.06028646 | 0.06991203 |
| Butanoate metabolism | 61 | 56 | 2.09161276 | 1.1361548 | 0.06134395 | 0.07033526 |
| Streptomycin biosynthesis | 12 | 11 | 2.06774132 | 1.13615673 | 0.06167077 | 0.07033526 |
| Valine, leucine and isoleucine biosynthesis | 15 | 15 | 2.1316621 | 1.13612152 | 0.07133274 | 0.08068786 |
| Zeatin biosynthesis | 3 | 1 | 2.22186941 | 1.13609808 | 0.07256124 | 0.08141017 |
| Fatty acid metabolism | 52 | 38 | 2.01047677 | 1.13615393 | 0.07396626 | 0.08231729 |
| Propanoate metabolism | 55 | 47 | 1.99682166 | 1.13615136 | 0.07582248 | 0.08370801 |
| Biosynthesis of vancomycin group antibiotics | 1 | 1 | 2.07213059 | 1.136114 | 0.09530025 | 0.10437646 |
| Biosynthesis of ansamycins | 1 | 1 | 2.0412 | 1.13610372 | 0.10045036 | 0.10915078 |
| Bisphenol degradation | 6 | 4 | 1.83627024 | 1.13606084 | 0.1306644 | 0.14087256 |
| Other types of O-glycan biosynthesis | 11 | 1 | 1.73995698 | 1.134569 | 0.13605017 | 0.14332003 |
| Glycosaminoglycan biosynthesis - keratan sulfate | 9 | 1 | 1.73995698 | 1.134569 | 0.13605017 | 0.14332003 |
| Glycosphingolipid biosynthesis - lacto and neolacto series | 24 | 1 | 1.73995698 | 1.134569 | 0.13605017 | 0.14332003 |
| Secondary bile acid biosynthesis | 1 | 1 | 1.85616217 | 1.13606051 | 0.13775027 | 0.14401164 |
| Terpenoid backbone biosynthesis | 23 | 20 | 1.72696922 | 1.13615187 | 0.1473256 | 0.15286416 |
| Biosynthesis of unsaturated fatty acids | 9 | 5 | 1.7317207 | 1.13611707 | 0.16383328 | 0.16872383 |
| D-Arginine and D-ornithine metabolism | 2 | 2 | 1.6073305 | 1.1361439 | 0.20445917 | 0.2090027 |
| Polyketide sugar unit biosynthesis | 4 | 4 | 1.55541929 | 1.13614467 | 0.21729733 | 0.22049288 |
| N-Glycan biosynthesis | 42 | 7 | 1.39351394 | 1.13579753 | 0.27576846 | 0.27778137 |
| Synthesis and degradation of ketone bodies | 5 | 5 | 0.88118043 | 1.13613234 | 0.58481416 | 0.58481416 |
