## Supplementary figures and images for "Response and oil degradation activities of a northeast Atlantic bacterial community to biogenic and synthetic surfactants"

### Supplementary Figure S1

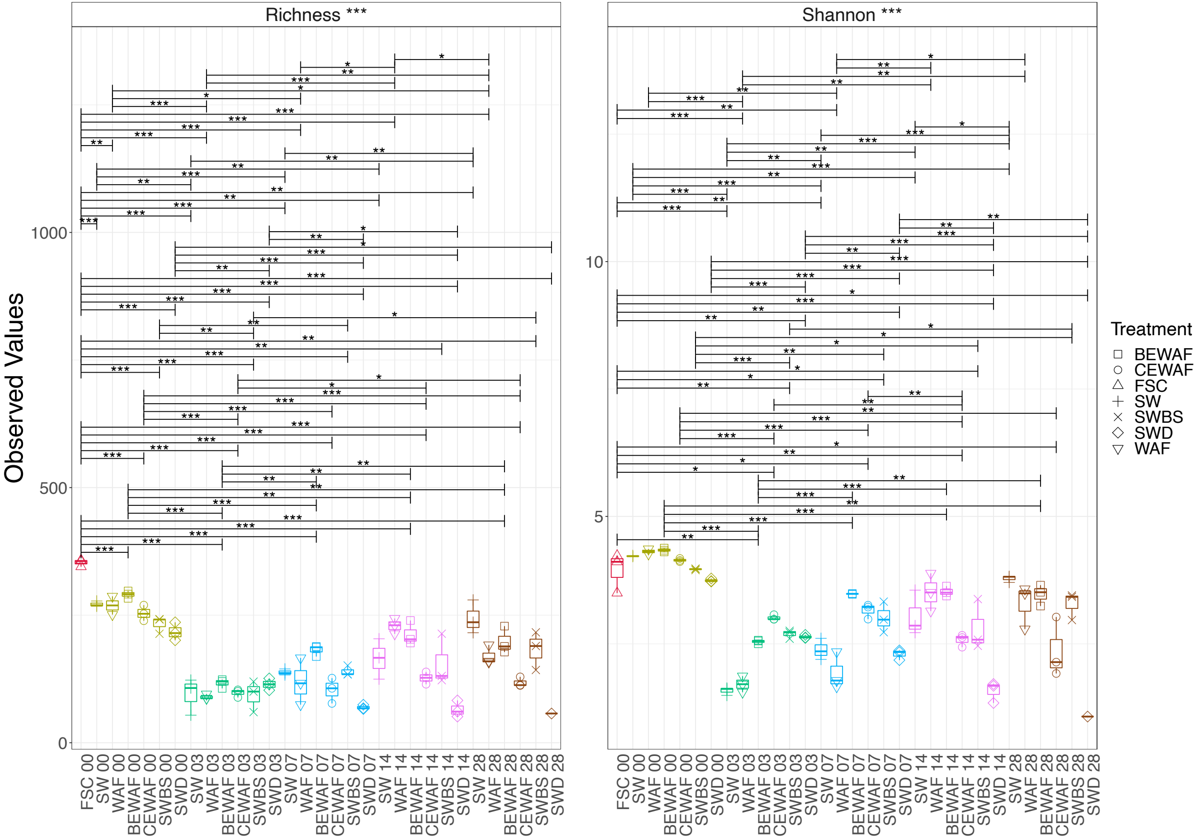

### Supplementary Figure S5

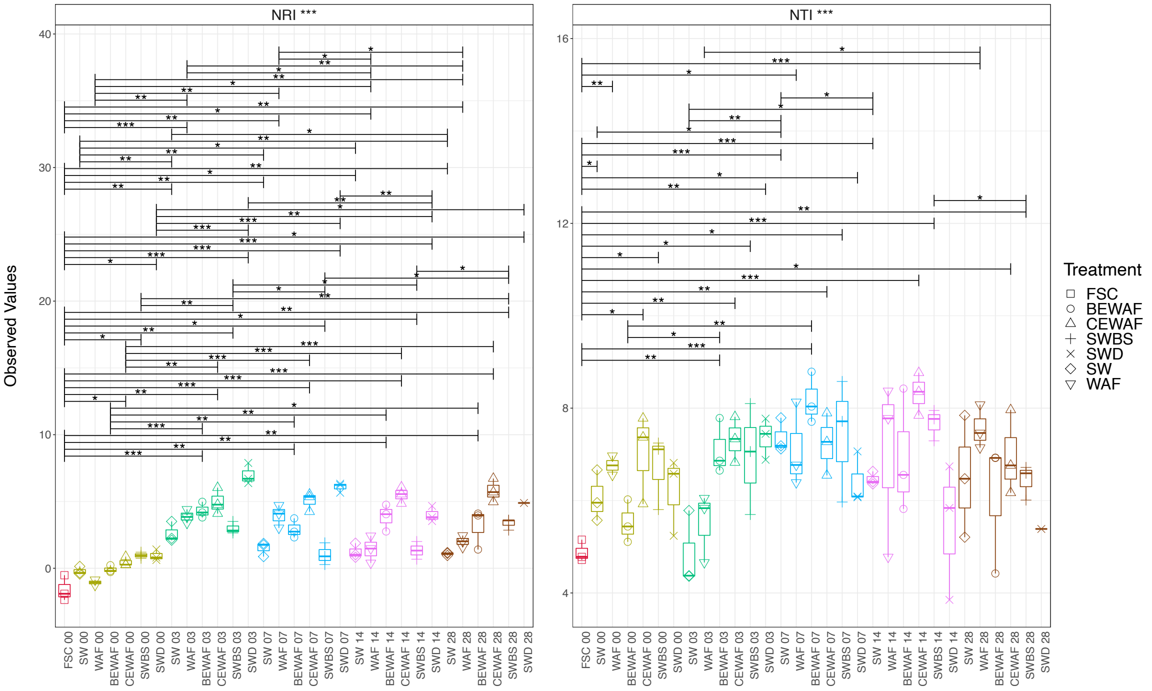

### Supplementary Figure S6

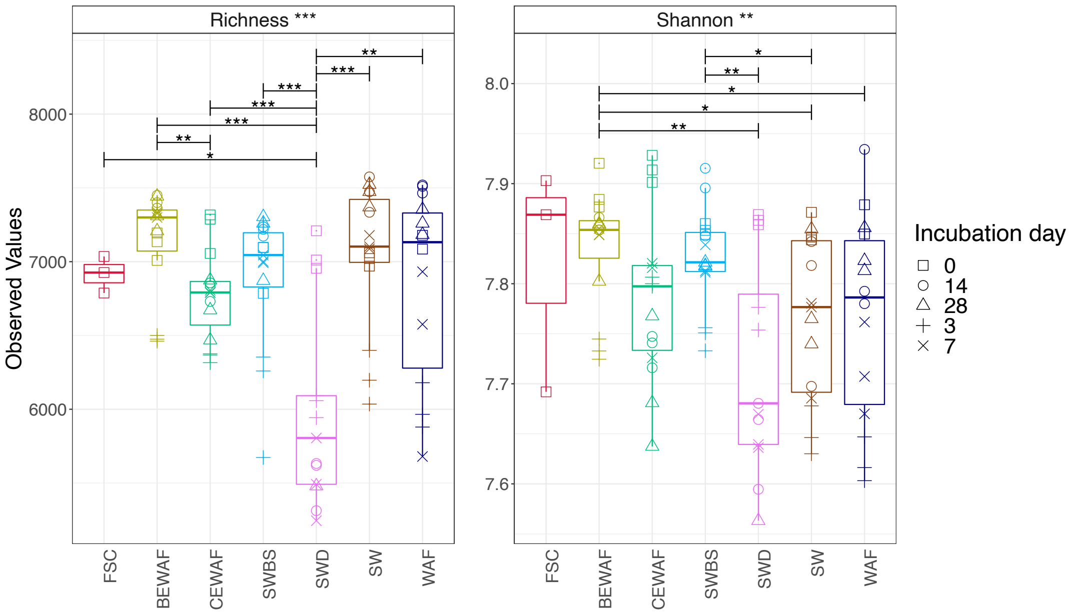

### Supplementary Figure S7

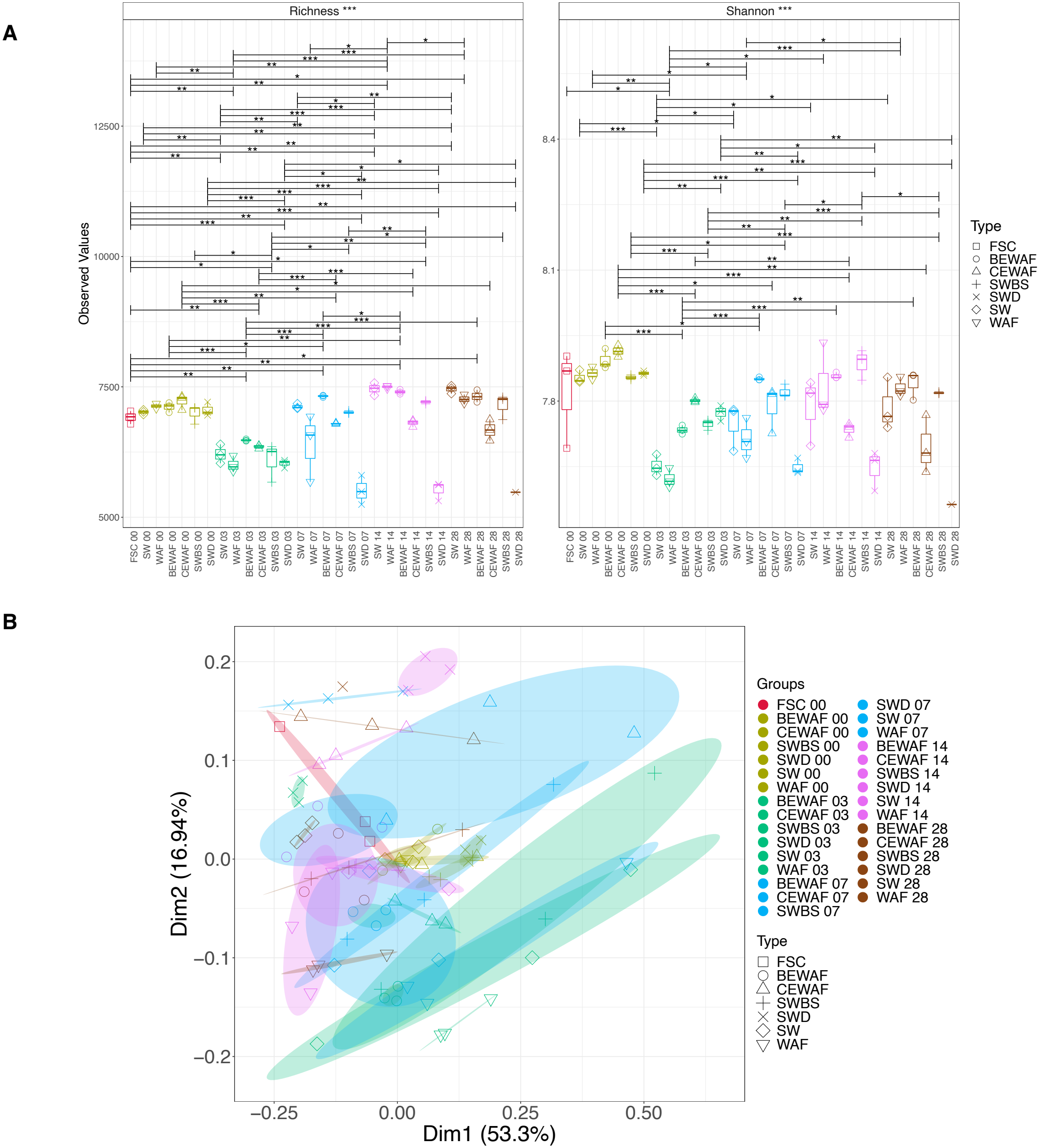

### Supplementary Figure S8

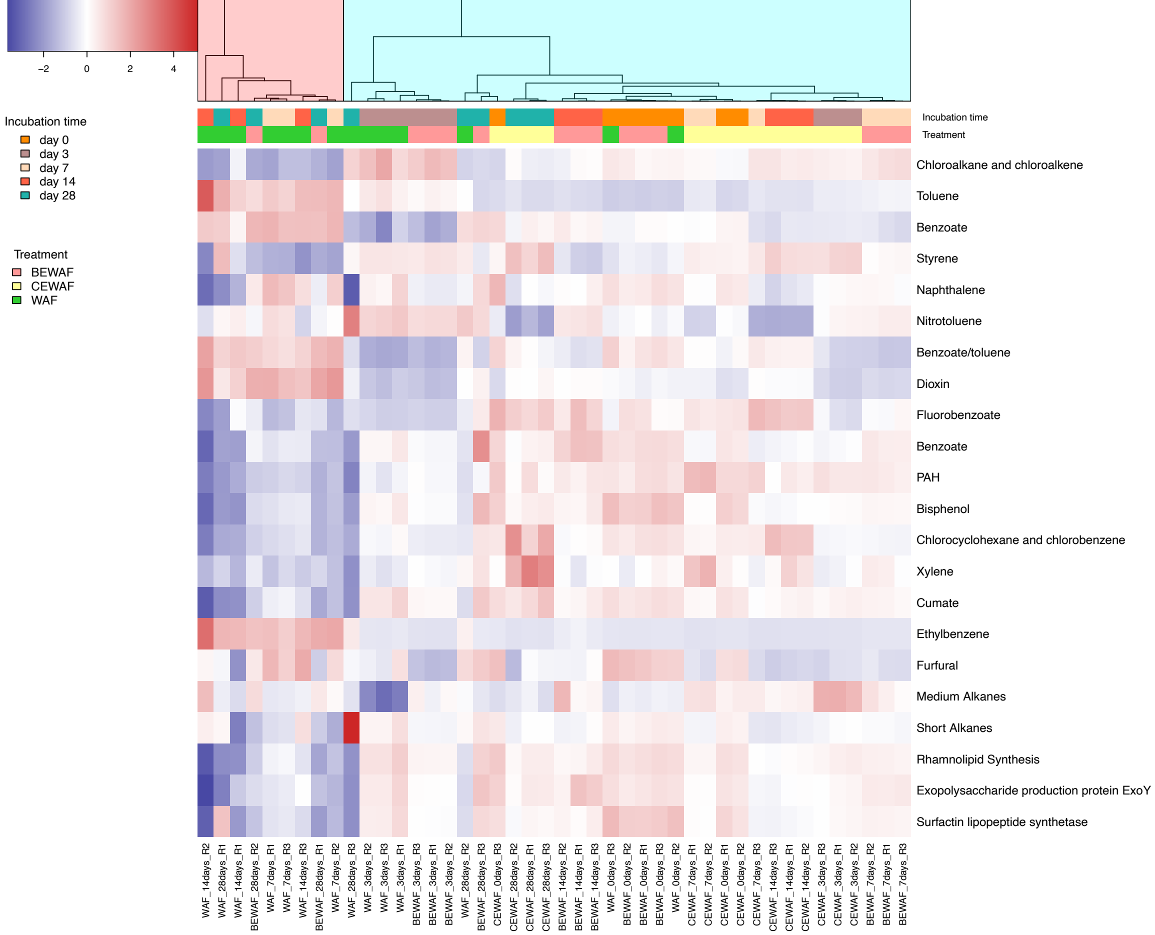

### Supplementary Figure S9

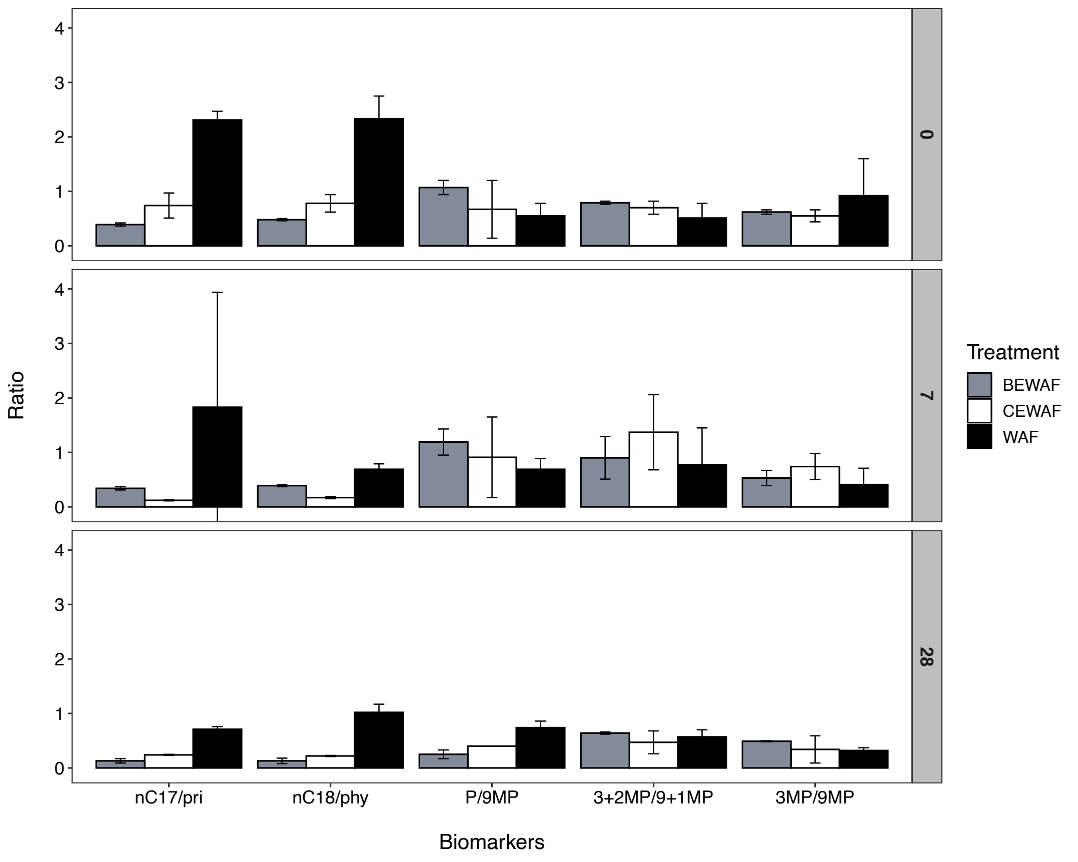
