## Supplementary Figure S4 for "Response and oil degradation activities of a northeast Atlantic bacterial community to biogenic and synthetic surfactants"

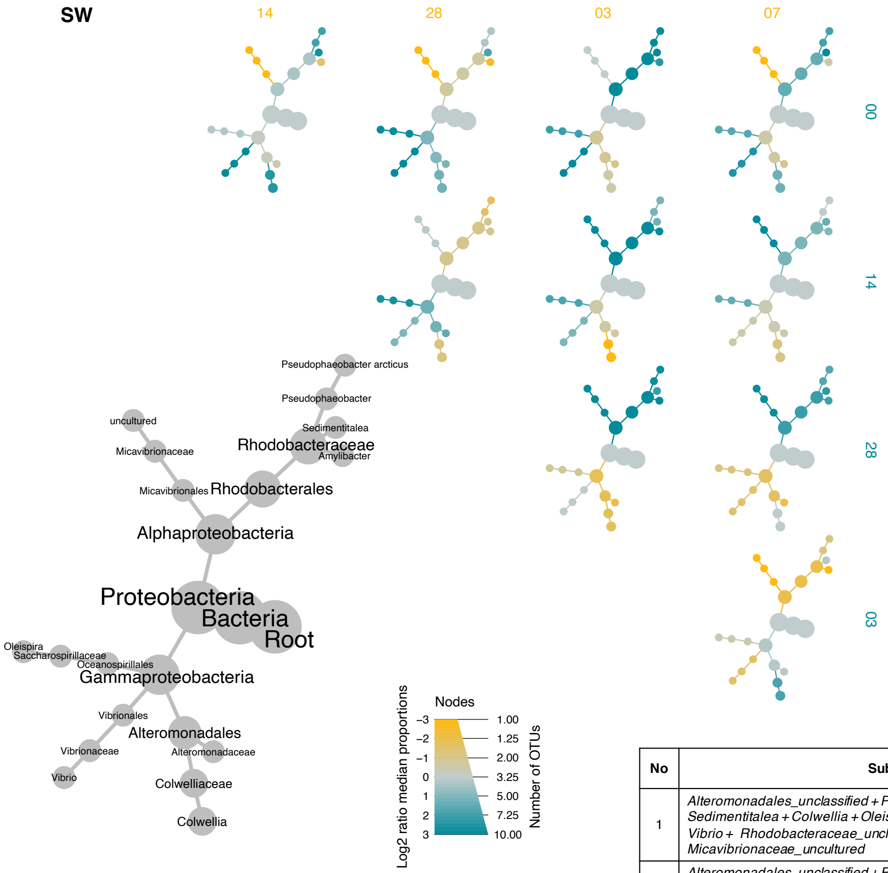


| No | Subset | Correlation with full ASV table | PERMANOVA |
|---|---|---|---|
| 1 | *Alteromonadales_unclassified + Pseudophaeobacter + Sedimentitalea + Colwellia + Oleispira + Amylibacter + Colwellia + Vibrio + Rhodobacteraceae_unclassified + Micavibrionaceae_uncultured* | 0.950 | $R^2 = 0.162$<br>P = 0.054 |
| 2 | *Alteromonadales_unclassified + Pseudophaeobacter + Sedimentitalea + Colwellia + Oleispira +Amylibacter + Colwellia + Vibrio + Rhodobacteraceae_unclassified* | 0.946 | $R^2 = 0.159$<br>P = 0.068 |
| 3 | *Alteromonadales_unclassified + Pseudophaeobacter + Sedimentitalea + Cowellia + Oleispira + Amylibacter + Vibrio + Rhodobacteraceae_unclassified* | 0.937 | $R^2 = 0.145$<br>P = 0.097 |
